# OoCount: A Machine-Learning Based Approach to Mouse Ovarian Follicle Counting and Classification

**DOI:** 10.1101/2024.05.13.593993

**Authors:** Lillian Folts, Anthony S. Martinez, Corey Bunce, Blanche Capel, Jennifer McKey

**Affiliations:** Section of Developmental Biology, Department of Pediatrics, University of Colorado Anschutz Medical Campus, Aurora, CO, USA; Konrad Lorenz Institute for Evolution and Cognition Research, Klosterneuburg, Austria; Department of Cell Biology, Duke University Medical Center, Durham, NC, USA

**Keywords:** Ovary, oocyte, mouse, machine learning, bioimage analysis, microscopy, follicle counting

## Abstract

The number and distribution of ovarian follicles in each growth stage provides a reliable readout of ovarian health and function. Leveraging techniques for three-dimensional (3D) imaging of ovaries *in toto* has the potential to uncover total, accurate ovarian follicle counts. However, because of the size and holistic nature of these images, counting oocytes is time consuming and difficult. The advent of deep-learning algorithms has allowed for the rapid development of ultra-fast, automated methods to analyze microscopy images. In recent years, these pipelines have become more user-friendly and accessible to non-specialists. We used these tools to create OoCount, a high-throughput, open-source method for automatic oocyte segmentation and classification from fluorescent 3D microscopy images of whole mouse ovaries using a deep-learning convolutional neural network (CNN) based approach. We developed a fast tissue-clearing and spinning disk confocal-based imaging protocol to obtain 3D images of whole mount perinatal and adult mouse ovaries. Fluorescently labeled oocytes from 3D images of ovaries were manually annotated in Napari to develop a machine learning training dataset. This dataset was used to retrain StarDist using a CNN within DL4MicEverywhere to automatically label all oocytes in the ovary. In a second phase, we utilize Accelerated Pixel and Object Classification, a Napari plugin, to classify labeled oocytes and sort them into growth stages. Here, we provide an end-to-end protocol for producing high-quality 3D images of the perinatal and adult mouse ovary, obtaining follicle counts and staging. We also demonstrate how to customize OoCount to fit images produced in any lab. Using OoCount, we can obtain accurate counts of oocytes in each growth stage in the perinatal and adult ovary, improving our ability to study ovarian function and fertility.

**Graphical Abstract:** 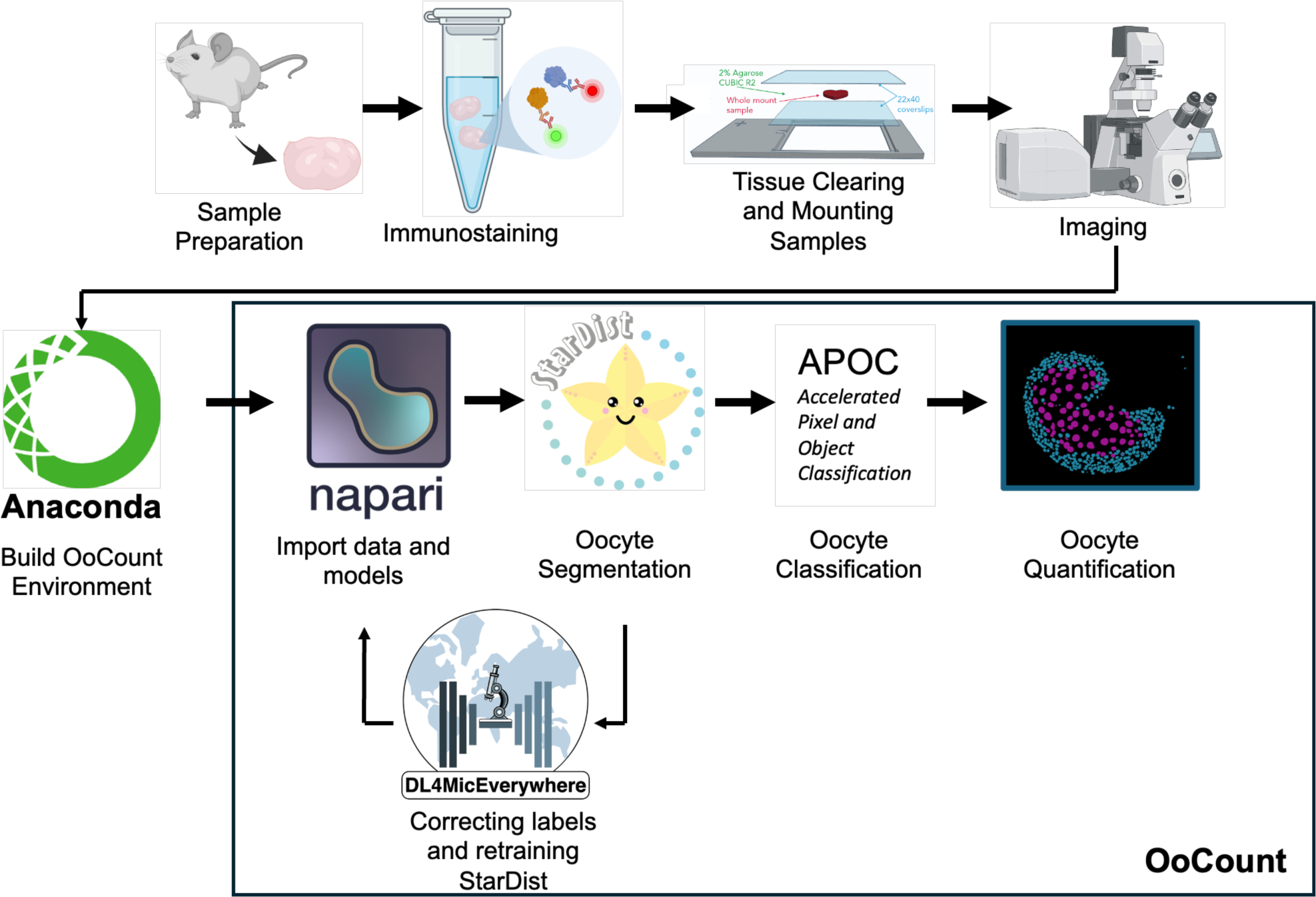

**Summary sentence:** This protocol introduces OoCount, a high-throughput, open-source method for automatic oocyte segmentation and classification from fluorescent 3D microscopy images of whole mouse ovaries using a machine learning-based approach.

## Introduction

Mammalian females are born with a finite supply of oocytes; thus, reproductive longevity is determined by the number of oocytes endowed at birth and the rate of depletion of this pool. Ovarian follicles, the functional units of the ovary, are composed of an oocyte surrounded by granulosa cells. During each reproductive cycle, a small number of primordial follicles, the quiescent follicles that make up the ovarian reserve, are activated and progress through growth stages. Eventually only a small subset of growing follicles in each cycle (or a single one in humans) will reach ovulation, while most others undergo atresia [1–3]. The number and distribution of ovarian follicles in each growth stage has provided the field of ovarian biology with a reliable readout of ovarian health and function [4,5]. This quantification is typically performed by extrapolating total follicle counts from histological sections of the ovary. However, due to the heterogeneous nature of ovarian tissue and variations in sectioning and extrapolation methods, reported follicle counts vary greatly between studies and are often inaccurate [5,6]. The recent development of techniques for three-dimensional (3D) imaging of ovaries *in toto* have opened up the possibility of obtaining robust, accurate, and near-absolute follicle counts [7–17].

3D imaging of ovaries *in toto* is providing the field with crucial spatial information on the regulation of follicle activation and growth, alongside improved follicle counts. However, extracting biological information from these large image datasets is challenging and time consuming. Fortunately, the advent of machine-learning algorithms allowed the rapid development of ultra-fast, automated methods to analyze large amounts of microscopy data [18–20]. In recent years, more and more groups have created user-friendly programming notebooks and code that provide biologists with accessible ways to utilize machine-learning in their research [21–27]. We leveraged tools including Napari [28], DL4MicEverywhere [22], StarDist [27] and Accelerated Pixel and Object Classification (APOC) [29] to create OoCount. We developed a fast tissue-clearing and spinning disk confocal-based imaging protocol to obtain 3D images of whole mount perinatal and adult mouse ovaries. We trained the existing StarDist convolutional neural network (CNN) to automatically label all oocytes in the perinatal and adult mouse ovary. In a second phase, we trained APOC, a machine-learning-based classifier to classify labeled oocytes and sort them into growth stages. With OoCount, users can obtain accurate counts of oocytes in each growth stage in the perinatal and adult ovary, improving our ability to study ovarian function and fertility.

### Overview of methods

The following protocol details the steps users can take to implement OoCount for analysis of follicle distribution in the perinatal or adult ovary. In steps 1-4, we provide a complete workflow for producing 3D images of the ovary compatible with our current version of OoCount. In steps 5-9, we detail the steps to utilize OoCount to segment and classify oocytes from 3D images of perinatal and adult ovaries. To segment (identify) oocytes we created custom StarDist models for perinatal and adult ovaries (https://www.mckeylab.com/oocount) [27]. To classify oocytes into growth stages, we provide instructions on training Accelerated Pixel and Object Classification (APOC) for perinatal and adult ovaries [29]. If the images generated in a different lab are incompatible, or do not yield satisfactory results with our version of OoCount, we have also included Step 9, which details the tools and approaches required to optimize OoCount for different types of images.

If using ovaries dissected from mice postnatal day (P) 8 or younger, we suggest following the “perinatal protocol” for tissue preparation and immunostaining. If using ovaries dissected from mice P9 and older, we suggest following the “adult protocol” for preparation and immunostaining. Our perinatal version of OoCount only distinguishes between growing and quiescent follicles, while our adult version of OoCount distinguishes primordial, primary, secondary, and antral follicles. This is important to note before using this workflow.

The entire pipeline was developed using samples from CD-1 mice (Charles River Laboratories). The use of animals to develop this method was conducted in accordance with protocols approved by the Institutional Animal Care and Use Committee of the University of Colorado Anschutz Medical Campus (IACUC protocol #1262) and Duke University (IACUC protocol #A089-20-04 9N).

In this manuscript, we provide protocols for the following steps:

1. *Sample preparation*. Ovaries are dissected from perinatal or adult mice, fixed and dehydrated before immunostaining.
2. *Immunostaining.* Ovaries are processed for fluorescent immunostaining to visualize oocytes. The reagents and antibodies required for this step are listed in Table 1 and Table S1.
3. *Tissue clearing and preparation for imaging.* Perinatal ovaries are embedded in a CUBIC-based clearing gel. Adult ovaries are cleared using Ethyl Cinnamate (ECi). In both cases, ovaries are mounted for imaging in 3D printed coverslip-holders, which we colloquially called “Coverslip’n’slide” (https://www.mckeylab.com/3d-prints).
4. *Imaging.* Mounted cleared ovaries are imaged using a spinning disk confocal microscope.
5. *Setting up the OoCount workflow and python environment*. Anaconda Navigator is downloaded onto the computer used for image analysis and a new python environment is created for OoCount.
6. *Segmenting oocytes with StarDist.* Using the OoCount-StarDist model, oocytes are segmented from the image.
7. *Classifying oocytes with APOC*. Oocytes are classified into growth stages using a marker of follicle activation.
8. *Obtaining total counts.* The total number of oocytes in each growth stage is quantified and displayed.
9. *Creating a custom StarDist model.* If the current OoCount model is not effective for images produced using different imaging setups or methods, we have provided the tools required to retrain OoCount.

**Table 1.** Summary of reagents required for staining and clearing perinatal and adult ovaries for OoCount analysis.

|  | Reagent | Component | Target Concentration | Catalog Number | Supplier | Preparation |
| --- | --- | --- | --- | --- | --- | --- |
| Perinatal Whole Mount Immunofluorescence | Methanol Gradient | Methanol<br>PBS | 75%, 50%, 25%<br>25%, 50%, 75% | 423950040<br>see below | Fisher Scientific<br>See Below | 40mL made in a 50 mL conical tube with 1X PBS as the diluent |
|  | 1X PBS | 10X PBS<br>Ultra-Pure Water | 1X PBS<br>Diluent | 70011069<br>10-977-023 | Gibco<br>Invitrogen | For a 500 mL solution make a 50mL aliquot of 10X PBS and bring to 500mL with diluent. Invert to mix. |
|  | PBS 1% Tx-100 | Triton X-100<br>1X PBS | 1%<br>Diluent | X100-500ML<br>70011069 | Sigma-Millipore<br>Gibco | Prepare a 10% Tx-100 solution as a working stock by mixing 10ml Triton X-100 with 90mL 1X PBS. Dilute to 1% by making a 5mL aliquot of 10% Tx-100 and bring to 50mL with 1X PBS. |
|  | PBS 0.1% Tx-100 | Triton X-100<br>1X PBS | 0.1%<br>Diluent | X100-500ML<br>70011069 | Sigma-Millipore<br>Gibco | Use 10% Tx-100 solution as a working stock. Then make a 0.1% solution by making a 500uL aliquot of 10% Tx-100 and bring to 50mL with 1X PBS. |
|  | Block Solution | Horse Serum<br>PBS 1% Tx-100 | 10%<br>Diluent | 26-050-088<br>See above | Gibco<br>See Above | Prepare a 10 mL solution by making a 1mL aliquot 100% horse serum and bring to 10mL with PBS 1% Tx-100. Invert to mix, store at 4°C for up to one week. |
|  | CUBIC R2 Solution | Urea<br>Sucrose<br>Triethanolamine<br>Hypure Water | 25%<br>50%<br>10%<br>x (g) to 100% | 501368682<br>501368582<br>90279-500ML<br>10-977-023 | Fisher Scientific<br>Fisher Scientific<br>Sigma-Millipore<br>Fisher Scientific | For a 100g solution combine: 25g urea, 50g sucrose and 10g triethanolamine then fill to 100g with hypure water. Scale up or down depending on use. Store at room temperature. |
|  | 2% CUBIC Agarose in CUBIC R2 | Agarose<br>R2 | 2%<br>Diluent | 50-255-228<br>See Above | Fisher Scientific<br>See Above | For a 100mL solution weigh 2g of agarose and add to a Erlenmeyer flask. Bring to 100mL with R2 reagent and heat gently until agarose goes into solution. Make 2mL aliquots and store at room temperature until needed. (Note: do not heat in a microwave, a heatblock with stir bar or 50mL in a falcon tube are recommended) |
| Adult Whole Mount Immunofluorescence | Isopropyl Alcohol | Isopropyl Alcohol<br>PBS | 75%, 50%, 25%<br>25%, 50%, 75% | 67-63-0<br>see below | Fisher Scientific<br>See Below | Make solutions using the combinations indicated. For example, for a 10mL solution of 75% isopropyl alcohol: add 7.5mL isopropanol to 2.5mL of 1X PBS. Do the same for the other solutions. |
|  | 1x PBS | 10X PBS<br>Ultra-Pure Water | 1X PBS | 70011069<br>10-977-023 | Gibco<br>Invitrogen | For a 500 mL solution make a 50mL aliquot of 10X PBS and bring to 500mL. Invert to mix. |
|  | PBS 0.2% Tx-100 | Triton Tx-100<br>1x PBS | 0.2%<br>Diluent | X100-500ML<br>70011069 | Sigma-Millipore<br>Gibco | Prepare a 10% Tx-100 solution as a working stock. Then make a 0.2% solution by making 1mL aliquot of 10% Tx-100 and bring to 50mL with 1X PBS. |
|  | Permeabilizing Solution | Glycine<br>PBS 0.2% Tx-100 | 2.3%<br>Diluent | G7126-500G<br>See Above | Sigma-Millipore<br>See Above | For a 50mL solution, add 1.15g glycine to a 50mL conical and bring to 50mL with PBS 0.2% Tx-100. Invert to mix. |
|  | Block Solution | Horse Serum<br>PBS 0.2% Tx-100 | 10%<br>Diluent | 26-050-088<br>See above | Gibco<br>See Above | For a 10mL solution make a 1mL aliquot of 100% horse serum and bring to 10mL with PBS 0.2% Tx-100. |
|  | Heparin Wash Solution (PTwH) | Heparin<br>PBS 0.2% Tx-100 | 10 ug/mL<br>Diluent | H3393-50KU<br>See Above | Sigma-Millipore<br>See Above | For a 10mL solution, make a 5mL aliquot of diluent and add 10uL of a 10mg/mL heparin stock solution. Bring to 10mL final volume and invert to mix. |
|  | Antibody Solution | Horse Serum<br>Heparin Wash Solution | 10%<br>Diluent | 26-050-088<br>See Above | Gibco<br>See Above | For a 10mL solution, make a 5mL aliquot of diluent and add 1mL 100% horse serum, fill to 10mL with diluent and invert to mix. |
|  | Ethyl Cinnamate | N/A | N/A | 112372-100G | Sigma Aldrich | Use as is. |

An overview of these protocol steps are presented in Figure 1. All reagents and solutions are listed in Table 1.

**Figure 1:**
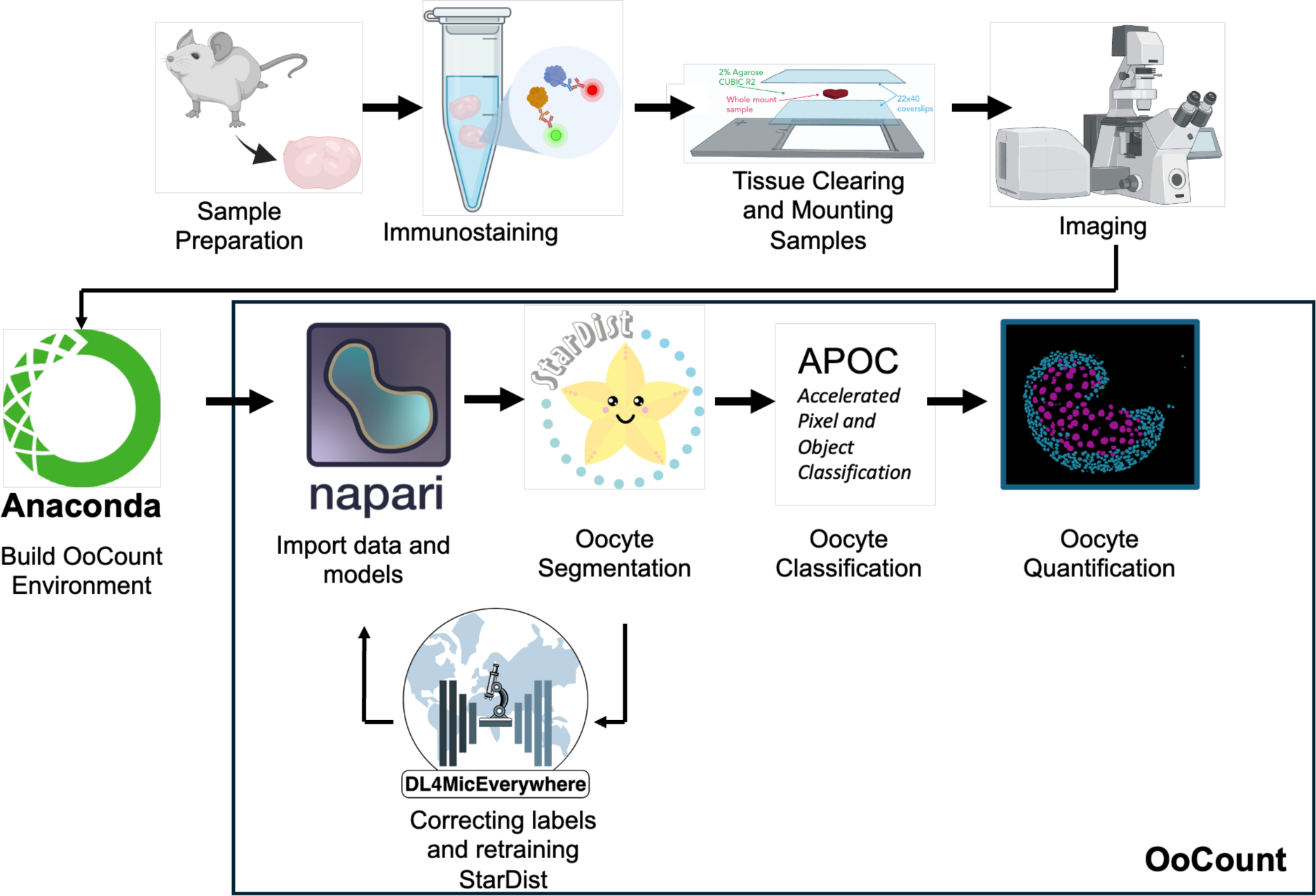
Overview of OoCount workflow. Schematic of the OoCount workflow including how to generate high quality 3D images of perinatal and adult ovaries to use within the computational pipeline.

All materials are listed in Table S2. All equipment and software used is listed in Table S3.

### Sample preparation

#### Perinatal (Pn) samples

Unless otherwise stated, use 500µl of solution at each step. For more detailed instructions on the dissection of ovarian tissue from perinatal mice, we suggest using the method outlined in the JoVE encyclopedia of experiments [30].

Pn.1.1 Euthanize female pups according to approved institutional regulations and dissect ovaries in 1X Phosphate Buffered Saline (PBS). Transfer the ovaries into a 1.5 mL tube with forceps.

Pn.1.2 Add 4% paraformaldehyde (PFA)/PBS to fix. Incubate for 30 minutes rocking at room temperature.

Pn.1.3 Following one quick rinse in 1X PBS, wash ovaries with 1X PBS for 20 minutes rocking at room temperature, or leave them in PBS at 4°C overnight without rocking.

Pn.1.4 Dehydrate ovaries in an increasing methanol (MeOH)/PBS gradient: 25%, 50%, 75%, 100% MeOH each for 15 minutes rocking at room temperature. Store ovaries in 100% methanol at −20°C until ready to use.

#### Adult (Ad) samples

Unless otherwise stated, use 1 ml of solution at each step. For more detailed instructions on the dissection of ovarian tissue from adult mice, we suggest using the method outlined by Converse and colleagues [31].

Ad.1.1 Euthanize female mice according to approved institutional regulations and dissect ovaries in PBS. Transfer them into a 1.5 mL tube with forceps. Ovaries should be removed from the bursa before further processing.

Ad.1.2 Add 4% PFA/PBS to fix. Incubate for 1 hour at room temperature rocking.

Ad.1.3 Following one quick rinse in 1X PBS, wash ovaries with 1X PBS for 20 minutes rocking at room temperature, or leave them in 1X PBS at 4°C overnight without rocking.

Ad.1.4 Dehydrate the ovaries in an increasing isopropanol (IPA)/PBS gradient: 30%, 50%, 70%, 100% IPA for 15 minutes each rocking at room temperature. Store ovaries in 100% IPA at −20°C until ready to use.

### Immunostaining

We recommend the following steps to perform whole-mount immunofluorescence staining on perinatal and adult mouse ovaries. We have provided detailed protocols in Supplementary protocols 1 (perinatal) and 2 (adult) and regularly updated versions can be found at https://www.mckeylab.com/protocols. In addition, we provide a list of solutions and reagents needed in Table S2. We used commercially available antibodies against DDX4 (raised in rabbit) to label oocytes, and against NR5A2 (raised in goat) to label granulosa cells (listed in Table S1).

**Note**: We serendipitously found that, in the perinatal ovary, the NR5A2 goat antibody labeled the nuclei of activated oocytes, in addition to labeling nuclei of activated granulosa cells. While NR5A2 is known to be expressed in the granulosa cells of active follicles [32–34], we have not confirmed that the NR5A2 protein is actually present in the nuclei of activated oocytes. Nonetheless, we have leveraged this unexpected staining as a marker of active oocytes in the perinatal ovary in our pipeline. If the user prefers a different marker of oocyte activation, such as GDF-9, we recommend using the rabbit GDF-9 antibody combined with the goat hVASA (human DDX4) antibody to label all oocytes (Table S1).

#### Perinatal samples

Start with samples that have been fixed and dehydrated according to Steps Pn1.1-1.4. Ovaries must be stored for at least one night in 100% methanol at −20°C. Unless otherwise stated, use 500µl of solution at each step. This section of the protocol takes a total of 3 days.

Pn.2.1: Rehydrate the samples in a decreasing MeOH/PBS gradient: 75%, 50%, 25% MeOH, 1X PBS for 15 minutes each rocking at room temperature.

Pn.2.2 Remove PBS and permeabilize for 20 minutes with PBS 0.1% Tx-100 rocking at room temperature.

Pn.2.3 Remove PBS 0.1% Tx-100 and incubate with freshly prepared block solution rocking at room temperature for 1 hour rocking.

Pn.2.4 Remove block solution and replace with primary antibody solution (1°AS). Primary antibodies should be diluted according to Table S2 in 250µl block solution. Incubate in 1°AS overnight at 4°C without rocking.

Pn.2.5 Remove 1°AS and wash samples 3 times for 20 minutes each with PBS 0.1% Tx-100 rocking at room temperature.

Pn.2.6 Remove PBS 0.1% Tx-100 and replace with secondary antibody solution (2°AS). For 2°AS, secondary antibodies should be diluted at 1:500 in 250µl of block solution and the solution should be filtered using a syringe-driven filter before adding to the samples. Place the tube with the samples in a dark box. Incubate in filtered 2°AS overnight at 4°C without rocking.

Pn.2.7 Remove 2°AS and wash samples 3 times 20 minutes with PBS 0.1% Tx-100 at room temperature rocking. Wash once with PBS for 15 minutes at room temperature rocking. Store samples at 4°C until ready to mount for imaging.

**Note**: For best results, we recommend mounting as soon as possible after staining. We have noted that the immunofluorescence signal is more stable in the mounting medium than in PBS, and mounted slides can be stored at 4°C in the dark for up to 5 days before imaging.

#### Adult samples

Start with samples that have been fixed and dehydrated according to Steps Ad1.1-1.4. Ovaries must be stored for at least one night in 100% IPA at −20°C. Unless otherwise stated, use 1ml of solution at each step. This section of the protocol takes 7-8 days.

Ad.2.1: Rehydrate the samples in a decreasing IPA/PBS gradient: 75%, 50%, 25% IPA for 20 minutes each at room temperature on a rocker. Wash for 20 minutes in PTx.2

Ad.2.2 Remove PTx.2 and replace with Permeabilizing Solution. Incubate overnight at 37°C without rocking.

Ad.Ad.2.3 Remove Permeabilizing Solution and incubate in freshly prepared Block solution at 37°C without rocking for 6 hours.

Ad.2.4 Remove Block solution and incubate with primary antibody solution (1°AS). Antibodies should be diluted according to Table S1 in 250µl block solution. Incubate in 1°AS for 3 nights at 37°C without rocking.

Ad.2.5 Remove 1°AS and wash samples 3 times 1 hour with heparin wash solution (PTwH) rocking at room temperature.

Ad.2.6 Remove PTwH and incubate with secondary antibody solution (2°AS). Antibodies should be diluted at 1:500 in 250µl block solution. Once prepared, filter 2°AS using a syringe-driven filter. Incubate in filtered 2°AS overnight at 4°C wihout rocking. After this step, the samples should be kept in the dark as much as possible.

Ad.2.7 Remove 2°AS and wash samples for 3 times 1 hour each with PTwH rocking at room temperature.

Ad.2.8 Remove PTwH and dehydrate samples using an increasing IPA/PBS gradient: 30%, 50%, 70%, 100% IPA for 20 minutes each rocking at room temperature. Proceed to tissue clearing (section Ad3) immediately after this step.

### Tissue clearing and mounting samples for imaging

#### Perinatal samples

Before performing this step, be sure to prepare 2ml aliquots of the CUBIC R2 agarose and have a 3D printed coverslip holder (coverslip’n’slide) prepared (Figure 2-1) (recipe for CUBIC R2 agarose can be found in Table 1; file for the coverslip’n’slide can be found at https://www.mckeylab.com/3d-prints) [35]. Materials are listed in Table S2. Detailed visual instructions for this step are illustrated in Figure 2.

**Figure 2:**
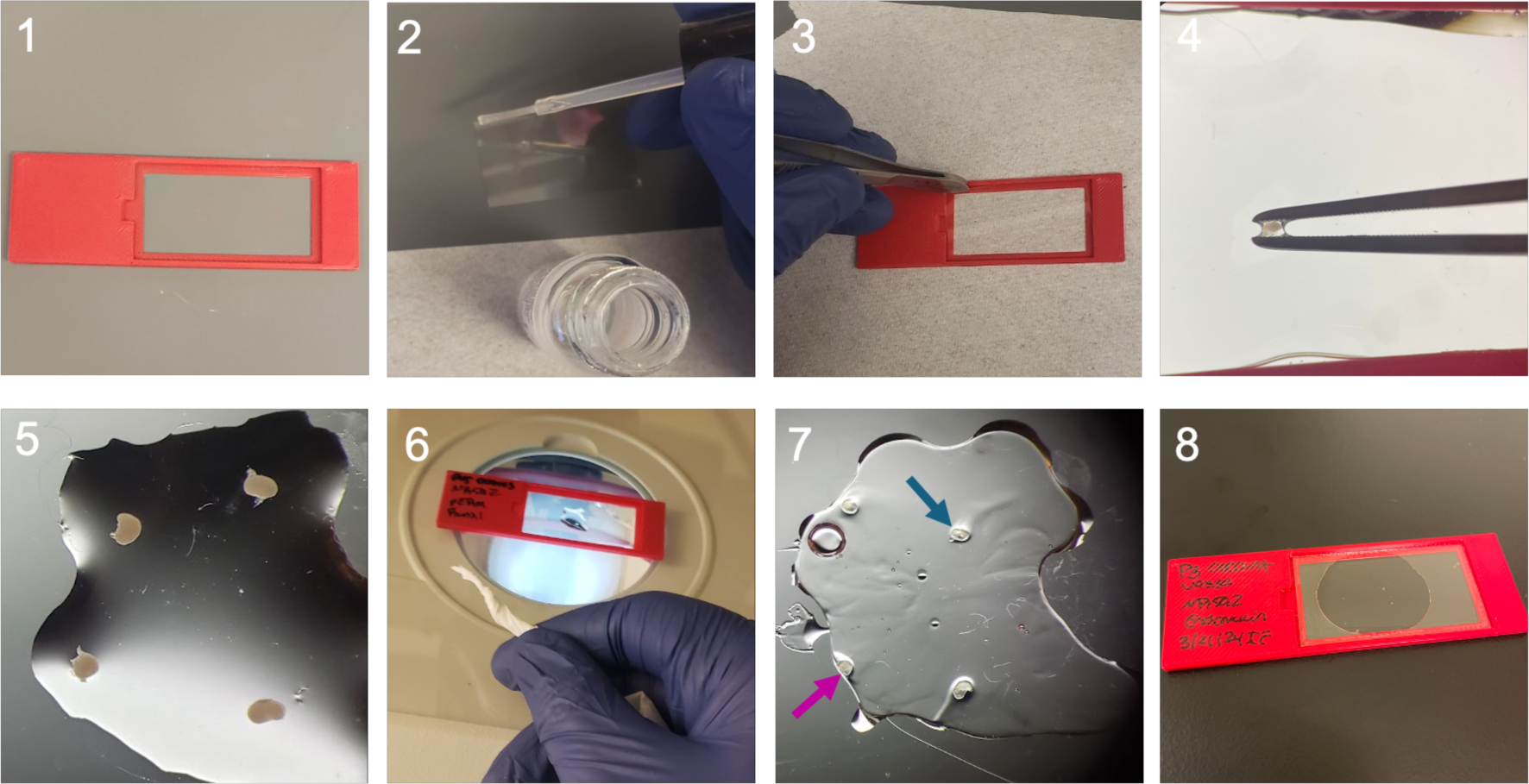
Mounting and preparing perinatal ovaries for imaging. 1. Print a coverslip’n’slide and gather supplies for mounting the samples. 2. Apply nail polish to the edges of a coverslip. 3. Attach the coverslip onto the coverslip’n’slide. 4. Using forceps, gently move the samples onto the coverslip. 5. Arrange the samples on the coverslip. 6. Using a tissue remove the excess liquid from the coverslip. 7. Slowly pipe the melted CUBIC R2 agarose around and in between the samples. Samples should be surrounded completely by agarose (as indicated by the blue arrow) and not partially covered (as indicated by the pink arrow) for best results. 8. Add a coverslip on top of the samples.

Pn.3.1 Place an aliquot of CUBIC R2 agarose at 95°C in a heat block 10 minutes before mounting the samples for clearing and imaging. **Note**: Avoid leaving the CUBIC R2 agarose at 95°C for too long, as it will burn and become yellow and slightly opaque.

Pn.3.2 Secure a coverslip onto the coverslip’n’slide by carefully lining the outer edges of the coverslip using clear coat nail polish. Do not use too little as this will make an imperfect seal, but not too much as this could refract the laser and make imaging your sample more difficult (Figure 2-2,3). Arrange the samples in the middle of the coverslip using forceps. Carefully remove any remaining PBS with tissue paper (Figure 2-4,5,6).

Pn.3.3 Using a pipet, slowly pipe approximately 100-200ul of the melted CUBIC R2 agarose around and in between the samples, being careful not to add the hot CUBIC R2 agarose solution directly onto the tissue. Eventually the pools of liquid CUBIC R2 agarose will come together to form a continuous blob with all tissue samples encased. Forceps can be used to gently nudge the agarose to cover the sample. Make sure that there is a good border of agar around your sample (Figure 2-7).

Pn.3.4 Place a coverslip over the samples and press lightly (Figure 2-8). **Note**: The amount of pressure applied on the coverslip will alter the 3D volume of the tissue. More pressure will decrease the time to acquire a Z-stack, but the rendered 3D images will look flatter.

#### Adult samples

Ethyl cinnamate (ECi) will be used to clear the adult ovaries. While ECi is not a hazardous material, inhalation of the vapors should be avoided. Materials are listed in Table S2. Detailed visual instructions for this step are illustrated in Figure 3.

**Figure 3:**
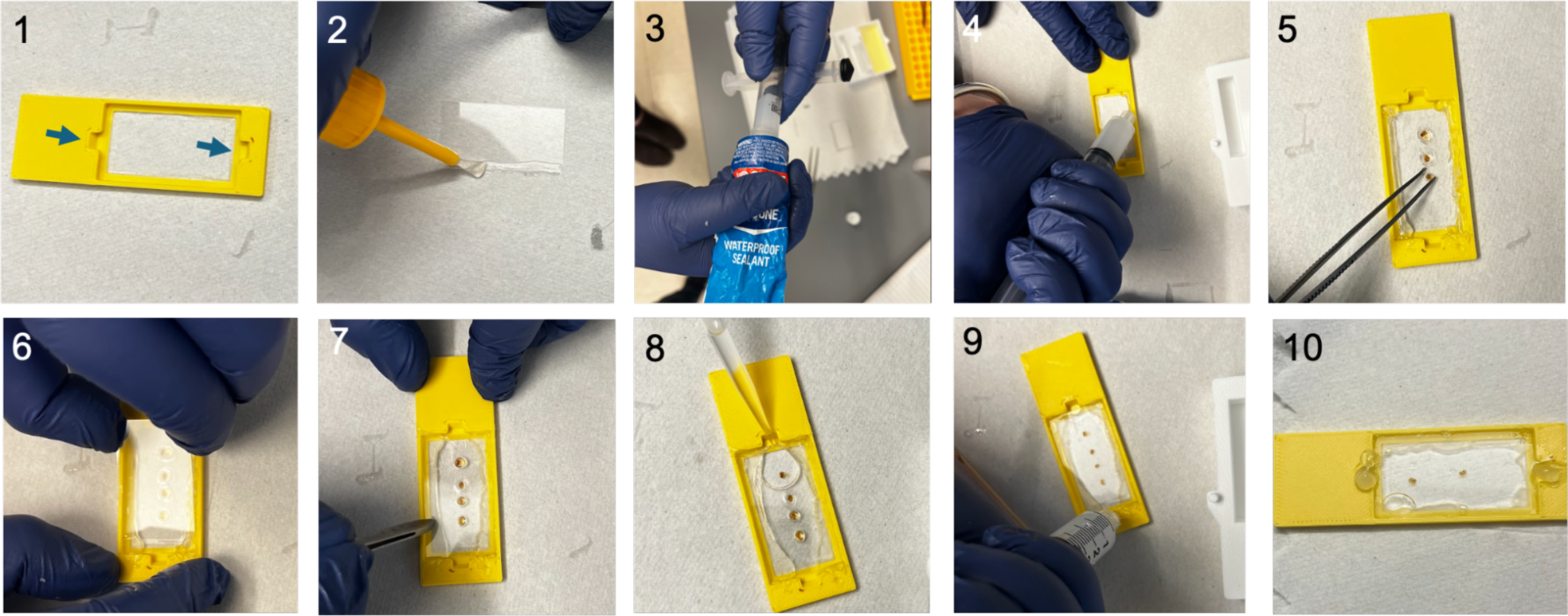
Mounting and preparing adult ovaries for imaging. 1. Print a coverslip’n’slide and gather materials for mounting the samples. Note the filling inlets on the coverslip’n’slide indicated with blue arrows. 2. Apply super glue around the edges of a coverslip and attach it to the coverslip’n’slide. 3. Load a syringe with sealant for more precise control over application. 4. Using the syringe, create a chamber using sealant by lining the edges of the coverslip. 5. Arrange the samples on the coverslip. 6. Apply a coverslip to the top of the sealant, creating a chamber. 7. Gently push on the top coverslip until it touches the samples. 8. Using a pipet, fill the chamber with ECi. 9. Plug the filling inlets with sealant. 10. Allow to cure for 1 hour before imaging.

Note that adult ovaries require a thicker coverslip chamber in the 3D printed coverslip holder (3mm for adult instead of 2mm for perinatal). The file for the 3mm coverslip’n’slide can be found at https://www.mckeylab.com/3d-prints. The thickness of the coverslip’n’slide coverslip chamber can be changed to fit the size of the tissue being imaged (modular coverslip slide file at https://www.mckeylab.com/3d-prints).

Ad.3.1 Replace 100% IPA with ECi, making sure to fill the entire tube or vial with ECi to avoid oxidation of the tissue, which prevents clearing. Invert the tube gently about 10 times, do not shake. Tube may be stored in the dark at room temperature until ready for imaging. Wait for the sample to be completely clear before proceeding to mounting and imaging (this can take a few hours).

Ad.3.2 Secure a base coverslip on the thick coverslip’n’slide using superglue. Attaching the coverslip with superglue is critical for mounting the ECi cleared samples. This creates a strong enough seal to prevent ECi from leaking out and damaging the objectives of the microscope and prevents skin contact with the ECi. Allow this to cure. Arrange the samples on the base coverslip using forceps (Figure 3-1,2).

Ad.3.3 Create a chamber using silicon or epoxy sealant along the edge of the base coverslip. Do not put sealant over the filling inlets (Figure 3-3). For precise control of the sealant, we recommend squeezing sealant into the back of a syringe (Figure 3-3,4) and using the syringe to apply the sealant to create a chamber. Transfer the samples onto the coverslip using forceps (Figure 3-5). Gently place a coverslip on top of the sealant making sure that the coverslip makes contact with the samples (Figure 3-6,7).

Ad.3.4 Fill the chamber with ECi with a pipet through the filling inlet. Fill the chamber as much as possible. Avoid creating bubbles over the samples (Figure 3-8).

Ad.3.5 Plug the inlets with sealant and allow 1 hour for the sealant to cure before imaging (Figure 3-9). Make sure that the chamber is well sealed and that ECi is not leaking by placing the slide on a clean tissue paper (Figure 3-10).

### Imaging

Imaging can be achieved on a variety of microscopes, as long as they are capable of imaging through large tissues (adult mouse ovaries are typically between 500µm and 1mm and perinatal mouse ovaries are about 200µm). Optimize microscope settings for fastest acquisition that allows for visualization of the smallest primordial follicles. We recommend a maximum Z interval of 5µm to ensure that primordial follicles are captured on at least 2 optical slices. To generate the images presented in this protocol, we used an Oxford Instruments Andor Dragonfly 202 Spinning Disk Confocal microscope with a 40μm disk pinhole (Oxford Instruments, Abingdon, UK), and a Leica DMi8 microscope stand (Leica Microsystems, Wetzlar, Germany) with an ASI Piezo stage controller (Applied Scientific Instrumentation, OR, USA). Images of perinatal or adult ovaries were acquired using a 25x water immersion (NA 0.95) or 10x dry (NA 0.45) objective (Leica Microsystems, Wetzlar, Germany), respectively. Images were captured by an Andor Zyla 4.2 plus sCMOS camera (Oxford Instruments Andor, Abingdon, UK). We used a high power laser engine (HLE) equipped with 637nm and 561nm lasers to visualize Alexa Fluor dye 647 (DDX4) and Cy3 (NR5A2), respectively. We used high laser powers (>10%) and low exposure times (<100ms) to optimize for fast acquisition. Perinatal or adult samples were imaged with a Z-interval of 1μm or 3μm, respectively. The tissue clearing achieved by following steps 1-4 will produce 3D images that allow for visualization through the entire perinatal or adult ovary (Figure 4, Videos 1 and 2). Note that this pipeline is not ideally suited for laser scanning confocal systems, as these will significantly increase acquisition time and sample photobleaching. We highly recommend using spinning disk or single plane illumination microscopy (SPIM / light sheet).

**Figure 4:**
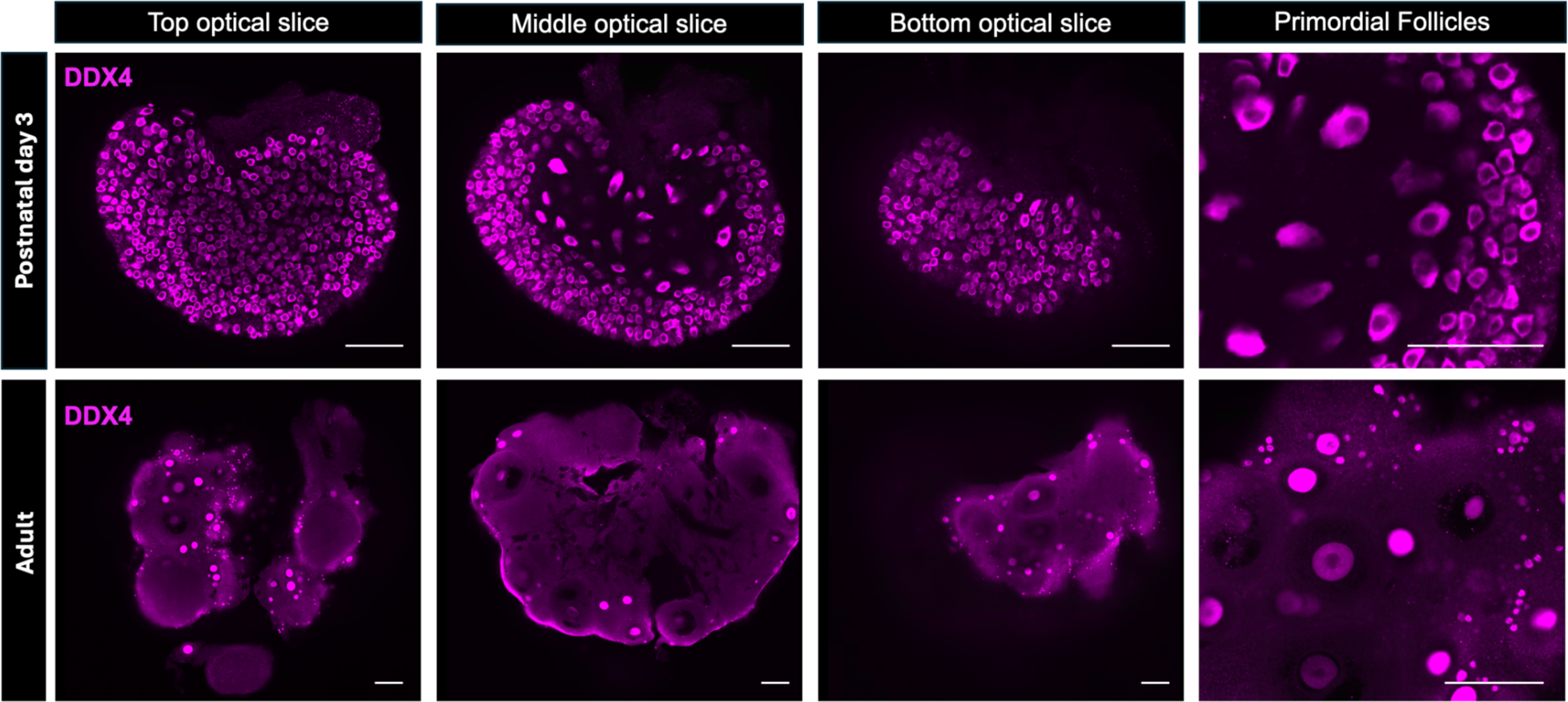
High resolution visualization of perinatal and adult ovaries *in toto*. Our immunostaining and clearing protocol is optimized for visualization of oocytes labeled with DDX4 antibodies (magenta) through the entire perinatal and adult ovary, including high resolution visualization of primordial follicles. Top optical slice is closest to the objective, while the bottom optical slice is furthest from the objective. The panel furthest to the right shows a zoom in on primordial follicles visualized in perinatal and adult ovaries. Scale bars: 150um.

### Setting up the OoCount workflow and Python environment

We recommend installing the required OoCount dependencies via Anaconda. Anaconda is a distribution of the Python programming language for scientific computing. Anaconda simplifies the installation of Python packages and the creation of “virtual environments”, which consist of a directory structure containing its own Python interpreter, standard library, and specific versions of additional dependencies required for a particular project. Anaconda provides a graphical user interface named “Anaconda Navigator” which can be used to create virtual environments and install dependencies.

Anaconda also provides a command-line tool named “conda” which can be used as an alternative to the graphical user interface.

The following instructions assume Anaconda and Anaconda Navigator have already been installed. Download Anaconda Navigator at https://www.anaconda.com/anaconda-navigator. **Note:** The computational portion of the pipeline is not different for analysis of adult and perinatal samples. Unless otherwise indicated, the steps will be the same.

5.1 Open Anaconda Navigator and click on Environments > Create to make a new environment for OoCount. Type in a name for the environment (e.g. OoCount) and select packages Python 3.9.7 then click “create”. Click the new environment in the side panel to activate it and click the “play” button to open the terminal.

**Note**: If you prefer not to use Anaconda Navigator, you can also create the conda environment directly in the terminal by running the following commands.

~~~
$ conda create -n OoCount -c conda-forge python=3.9
$ conda activate OoCount
~~~

This requires conda to be installed on your machine, installation instructions can be found at https://conda.io/projects/conda/en/latest/user-guide/install/index.html.

5.2 Once your OoCount environment is installed and activated, enter the following commands, one by one in the terminal:

~~~
$ pip install tensorflow==2.12
$ pip install matplotlib==3.8.2
$ pip install tifffile==2023.9.26
$ pip install tqdm==4.66.1
$ pip install imageio==2.33.0
$ pip install numba==0.58.1
$ pip install scikit-image==0.22.0
$ pip install napari==0.4.18
$ pip install stardist-napari==2022.12.6
$ pip install napari-accelerated-pixel-and-object-classification==0.14.1
$ pip install devbio-napari==0.10.1
~~~

This will install the specific versions of the packages used for OoCount into your conda environment.

**Note**: if using a PC with CUDA-compatible GPU and you want to use GPU-acceleration, you will need to install CUDAtoolkit:

~~~
$ pip install cudatoolkit=10.2
~~~

5.3 After installation, re-open Anaconda Navigator. Click on Environments > OoCount > open with terminal (Video 3). When the terminal opens, type ‘napari’ and a Napari window will open. Napari (Napari contributors, 2019) is a free and open-sourced python-based interactive viewer for multi-dimensional images. Napari can be used for state-of-the-art computational image analysis tasks comparable to FiJi [36] or Imaris (Bitplane, Inc).

5.4 To import images into Napari, the file can be dragged in or uploaded by clicking on File > Open. Import images as single channel tiffs or import multichannel tiffs and split channels in Napari. Once opened, the screen should resemble Figure 5. A brief overview of Napari can be seen in Video 4 but further instructions and tutorials on how to use the Napari viewer can be found at https://napari.org/stable/index.html.

**Figure 5:**
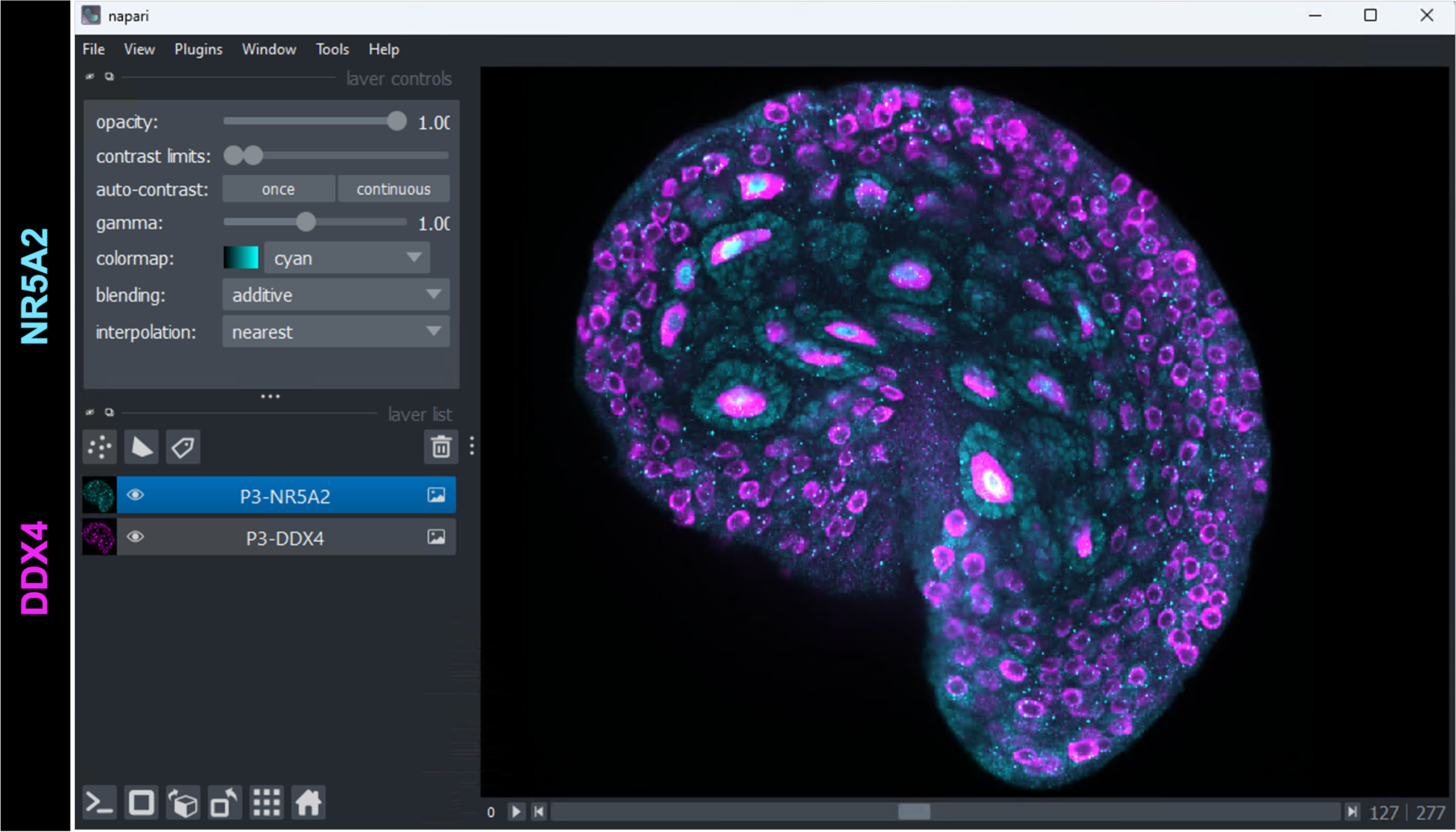
Opening images in Napari. Example screenshot of a multichannel image open in Napari. In a mouse perinatal ovary, DDX4 labels all oocytes and NR5A2 labels the nuclei of active oocytes and growing granulosa cells.

5.5 Before continuing, download the OoCount deliverables from our lab website (https://mckeylab.com/oocount). This folder contains the necessary files and code needed for this workflow, in addition to example data.

### StarDist-OoCount model to segment mouse oocytes

Before beginning this section, it is important to set up an organizational system to save images and labels. Create a new folder and create two subfolders, one named “Images’’ and the other named “Masks.” The “Images” folder should contain all the images you wish to segment, and the “Masks” folder is where you will later save the labels layers after running StarDist-OoCount.

To obtain a near-absolute count of oocytes in a 3D image of the ovary, the oocytes must be segmented from the image. In image analysis, object segmentation is the process of identifying specific objects or regions of interest within an image. StarDist is a segmentation tool that can run as a plugin in Napari (StarDist-Napari plugin) [27]. While it was originally created to segment fluorescent nuclei, we retrained the “3D_demo” StarDist model to segment oocytes from 3D images of perinatal and adult ovaries. We created two retrained StarDist models called StarDist-OoCount_Perinatal and StarDist-OoCount_Adult. To assess the efficacy of the training, we used training validation and training loss plots automatically rendered by the DL4MicEverywhere StarDist 3D training notebook (Figure S1A) [22,23]. In addition, we followed the quality control prompts of the training notebook, which perform comparisons between ground truth annotations and model prediction (Figure S1B). These comparisons yielded average Intersection over Union (IoU) values of 0.77 for OoCount_Perinatal, and 0.71 for OoCount_Adult. It is worth noting that the IoU is calculated as a pixel match between ground truth and prediction, so if the shape of the predicted label doesn’t match the shape of the manual annotation, the IoU decreases, even though the oocyte was accurately recognized as an object to label. Thus, the IoU is a conservative estimate of the efficacy of our StarDist-OoCount models.

After running StarDist-OoCount on an image in Napari, an output layer with labels will be created. The labels layer of the image contains the segmented oocytes.

6.1 Once an image with DDX4-labeled oocytes is imported into Napari, click Plugins > Napari-StarDist. A StarDist panel should open on the right side of the Napari window.

6.2 Keep all the default settings, except for those changed in Video 5. The Image Axis, Model Type, Custom Model, and Number of Tiles will need to be changed. *For perinatal ovaries*: Image Axis is ZXY, Model Type is Custom 2D/3D Model, Custom Model is FILE NAME, and Number of Tiles is 1,6,6 (Video 5). *For adult ovaries*: Image Axis is ZXY, Model Type is Custom 2D/3D Model, Custom Model is FILE NAME, and Number of Tiles is 1,10,10 (Video 5). After these settings have been adjusted, click Run. **Note**: The “number of tiles” setting is a z, y, x vector, and will need to be adjusted based on computational power and the size of the image to segment. If StarDist crashes when using these suggested values, consider increasing the number of tiles (e.g. 2,12,12) to reduce the computational load for StarDist. The *scaling* setting should be changed if the scale of the image to segment is significantly different from those that were used to train StarDist. For our retrained models, the voxel size for the perinatal training images was z=0.996μm, y=0.603μm, x=0.603μm and for the adult images, it was z=3μm, y=0.241μm, x=0.241μm. For optimal segmentation accuracy, the image to segment must be scaled to match the voxel size of the training data. The scaling setting takes a (z,y,x) vector. For example, if an image has a voxel size of z=0.996μm, y=0.241μm, x=0.241μm, x and y values of the image should be divided by 2 to match the StarDist-OoCount_perinatal model. In this case, the scaling setting should be set to (1, 0.5, 0.5). **Note**: the adult and perinatal models that we generated differ in scaling because perinatal images were captured with a 25x objective, while adult images were captured with a 10x objective. We recommend you use the model that is closest to the scale of the image to segment, as it may provide better results than the “age-matched” model.

Once the image has been segmented by OoCount-StarDist, it should resemble the output in Figure 6 and Video 5, where each oocyte is segmented with a different colored label. If there are missegmented oocytes, these can be corrected by following the steps in Video 6. Save the labels layer in the “Masks’’ folder with the **same file name** as the image.

**Figure 6:**
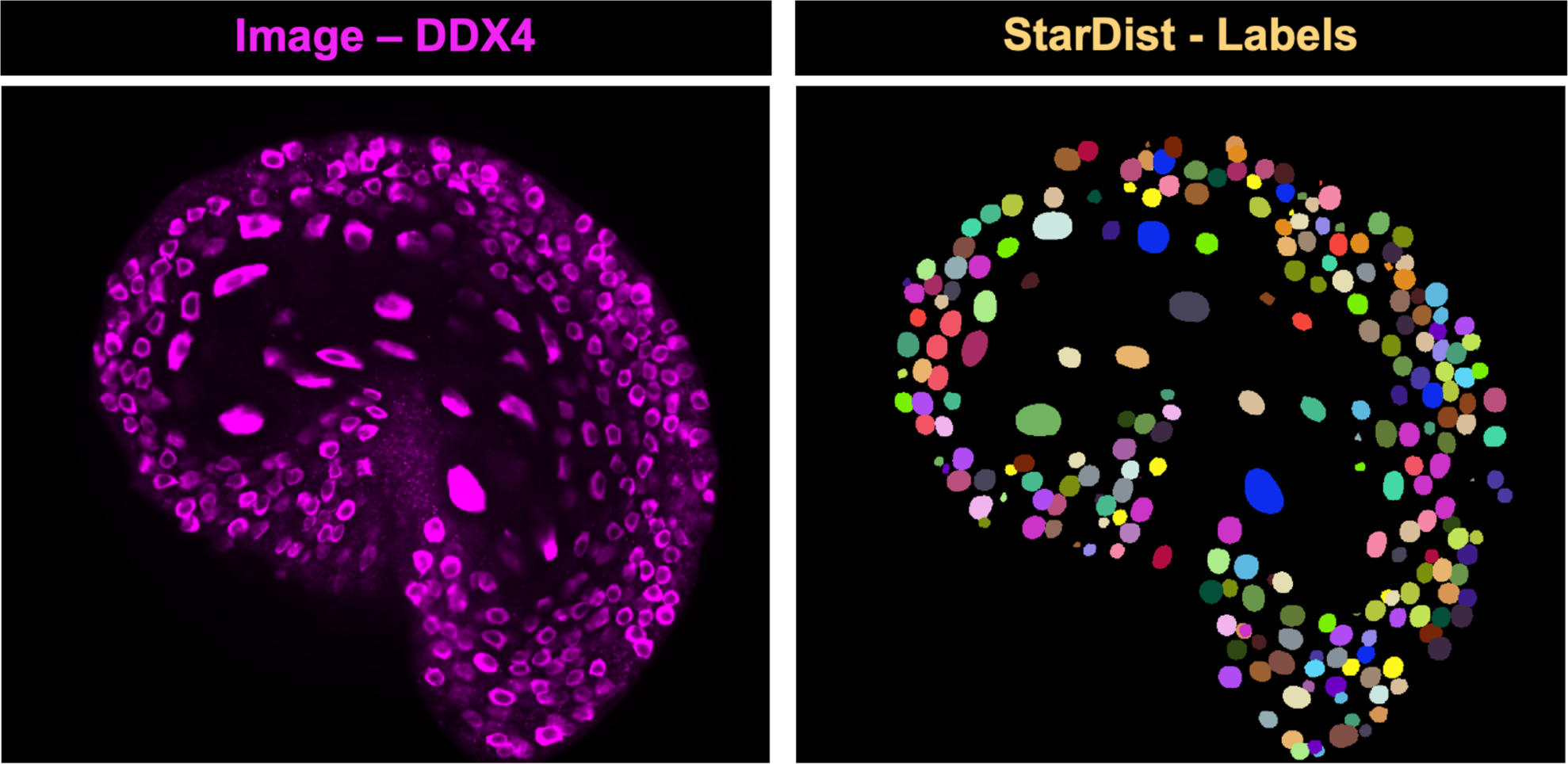
Segmenting oocytes from 3D images of ovaries. The StarDist-OoCount model segments oocytes (labeled using DDX4) from 3D images of perinatal and adult (not shown here) ovaries.

### Training APOC to classify oocytes

Once oocytes have been segmented from the image, they can be classified based on their growth stage. When performing object classification, objects within an image are assigned to certain classes or categories based on their features. Accelerated Pixel and Object Classifier (APOC) [29] is a classifier plugin for Napari [37]. APOC uses machine learning to train based on user-input, making it easy-to-use and versatile. Based on factors such as the expression of the activation marker NR5A2, pixel number, and signal intensity, APOC will classify oocyte labels created by StarDist-OoCount into growth stages with minimal training. For perinatal ovaries, oocytes are classified into one of two states: quiescent or active. In adult ovaries, we demonstrate how oocytes can be classified into one of four growth stages: primordial, primary, secondary, and antral. This analysis uses the segmentation result from StarDist-OoCount (mask file) and the individual channel images for the marker that will be used for classification (image file). In our case, we use DDX4 as a marker to segment all oocytes, and NR5A2 as a marker of follicle activation for classification (more specific guidance on follicle classification can be found in Figure S2). Any specific marker that distinguishes active from quiescent follicles should work. Alternatively, classification can be done for features other than growing or quiescent. For example, users may opt to use a cell death marker to quantify attrition or use expression of a protein of interest to quantify how many oocytes are expressing it. For additional information and support on using APOC in Napari, we recommend reading the user guide (https://github.com/haesleinhuepf/napari-accelerated-pixel-and-object-classification).

7.1 In Napari, open an image with the NR5A2 channel and the corresponding mask from StarDist (Figure 7 and Video 7). In Napari, click Plugins > Napari-accelerated-pixel-and-object-classification > object classification. An APOC panel should appear on the right side of the screen.

**Figure 7:**
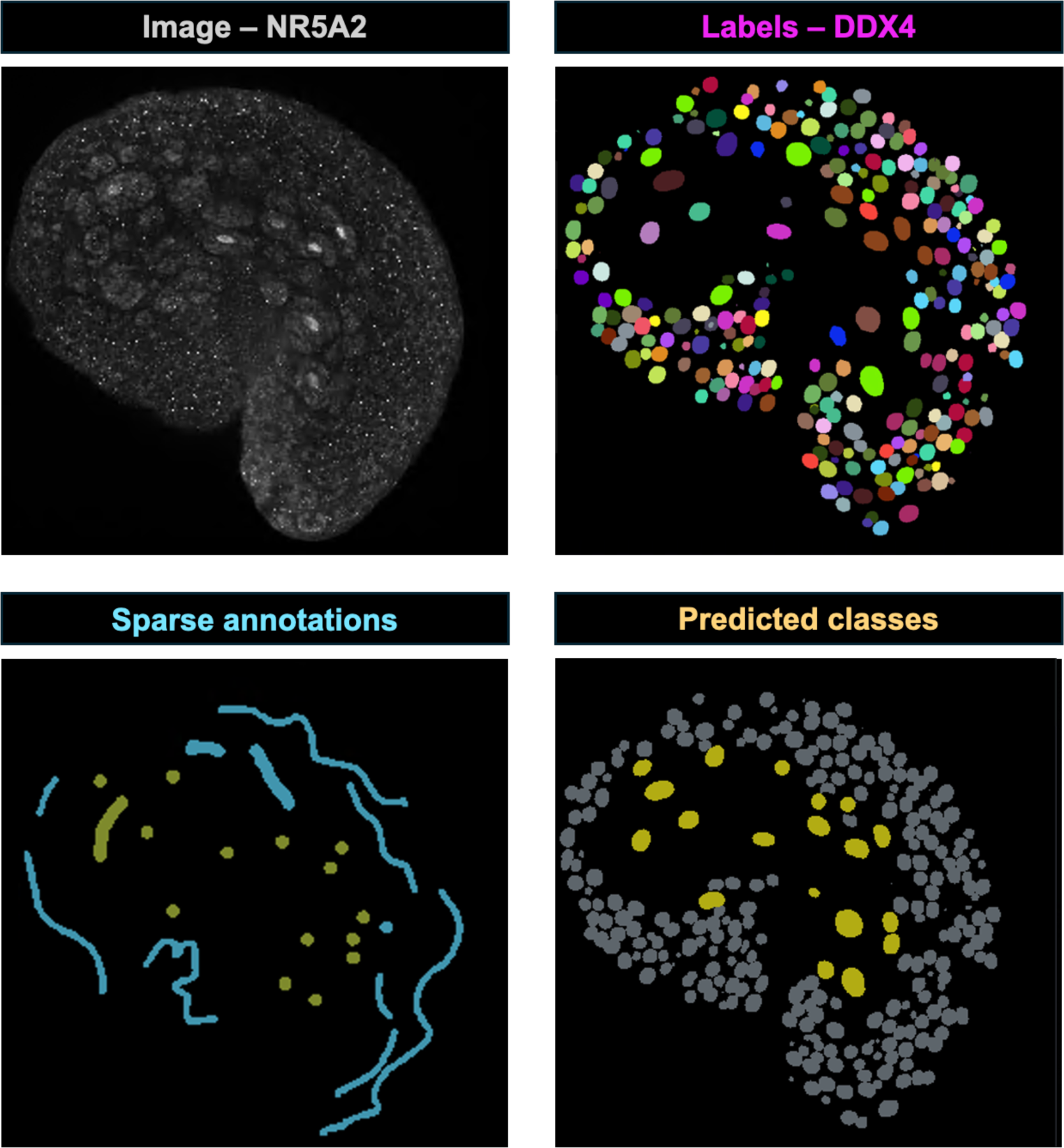
APOC classification in perinatal ovaries. When doing classification in APOC, the NR5A2 image (seen in top left panel) overlayed with the DDX4 labels (seen in top right panel) will be useful for determining activation of oocytes to assist in sparse annotations (seen in bottom left panel). After training, APOC classifies quiescent follicles (seen in the bottom right panel in grey) and active follicles (seen in bottom right panel in yellow) in the perinatal mouse ovary.

7.2 Create a new labels layer, and name it “annotation”. This new layer will hold your annotations for each class. Use the paintbrush tool to paint over the StarDist mask (Figure 7 and Video 7). This will assign a class number to each label based on the growth stage. For example, in perinatal ovaries, use label #1 for quiescent follicles and label #2 for active follicles. APOC works with sparse annotation, so there is no need to paint over every oocyte. Be sure to move through the Z-stacks and annotate oocytes at different depths and in different regions of the ovary. Repeat this for each class. **Note**: On the StarDist labels layers, change the “contour” value to 1 or 2 to toggle contour view instead of fill. This will allow you to see the underlying NR5A2 staining.

7.3 Once oocytes of all classes have been sparsely annotated, APOC can be run (Video 8). Be sure to save the classifier file where it is accessible for future use. Set the tree depth to 5 and the number of trees to 100. We obtain satisfactory classification results by selecting the following features for classification: Mean intensity, pixel count, standard deviation intensity, shape, centroid position, touching neighbor count, and average distance to touching neighbors. However, we recommend testing different combinations of features depending on your classification goals. Click Train.

7.4 Inspect the classification for errors. Using the same labels as 7.2, paint over errors in the annotation layer. Do not correct every erroneous label, so that APOC can attempt to correct them during retraining. Click Train again. Once APOC is achieving satisfactory classification, save the final APOC prediction output layer. **Note**: APOC will likely not classify oocytes with 100% accuracy but this can be manually corrected. Reopen the classification layer in Napari over the image with the NR5A2 channel and StarDist labels. The classification layer can now be edited using the label eraser and brush tools to correct any mis-classified oocytes (Video 9).

Once APOC has classified the oocytes, the output should resemble Figure 7 for perinatal ovaries, and Figure 8 for adult images. Classifier files created in APOC can reliably be used for all images within a single imaging batch, however, APOC may need to be retrained for new batches of images.

**Figure 8:**
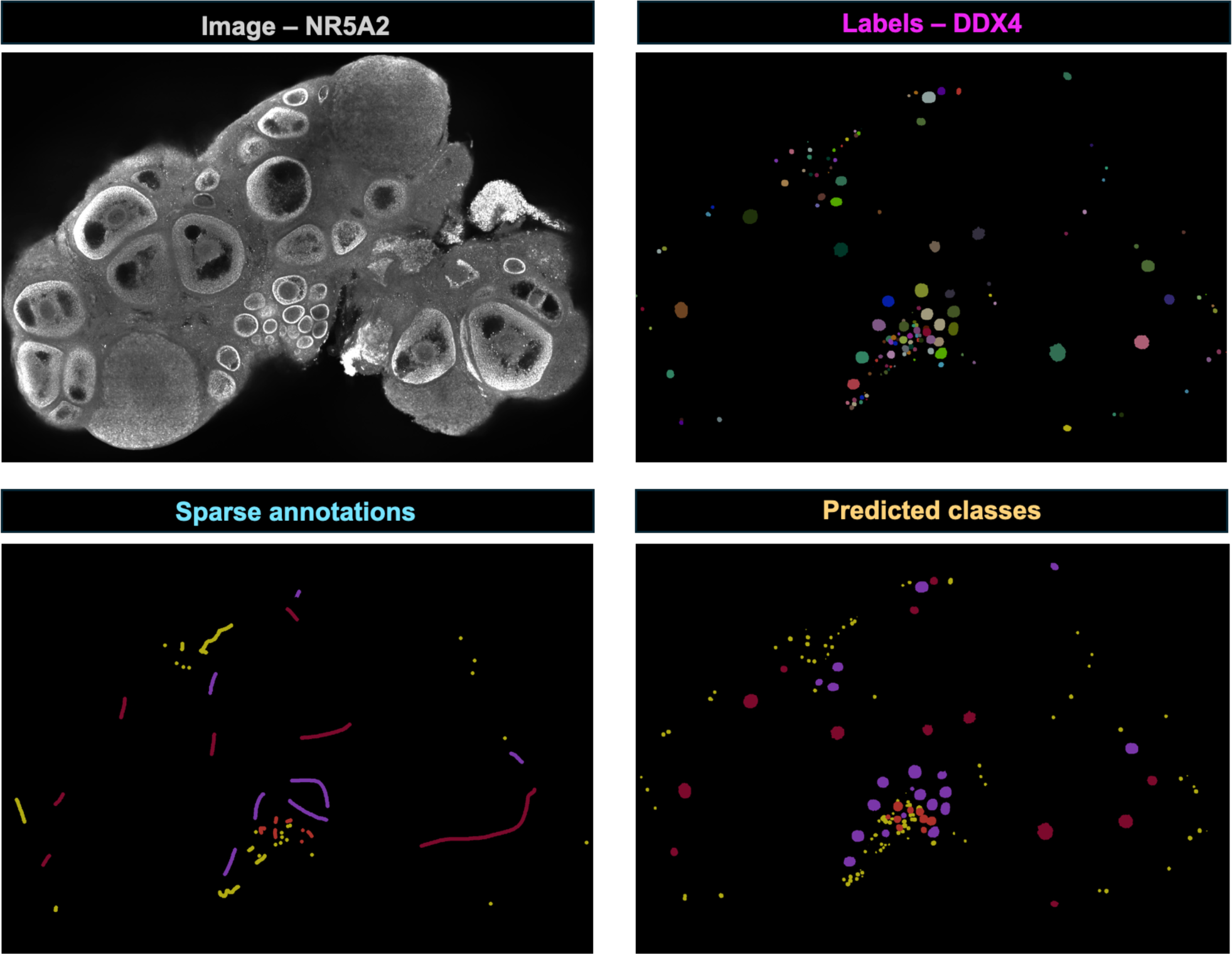
APOC classification in adult ovaries. When classifying follicles from the adult ovary, we recommend using the NR5A2 image layer (seen in the top left panel) overlayed with the DDX labels (seen in the top right panel) to determine the stage of the follicles. Use this to sparsely annotate the labels (seen in the bottom left panel). After training, APOC will classify follicles into primordial (yellow), primary (orange), secondary (purple), and antral (red).

### Getting total counts

To produce final oocyte counts, we used the Napari-gpt plugin (https://github.com/royerlab/napari-chatgpt) to generate code that would output oocyte counts by class [38]. With this code, for each unique label produced by StarDist, the label number assigned to it by APOC is determined and counted (e.g. 1 for quiescent and 2 for active).

8.1 Once the segmented oocytes from StarDist are classified into growth stages by APOC, open the file “Napari-OoCountResult.py” in any text editor (we use Visual Studio Code, Microsoft Corporation). Copy the code provided and paste it into the Napari console (Video 10). Click enter to run the code. Follow the prompts to obtain an output with the total number of oocytes and the number of oocytes in each growth stage (Figure 9). The output is generated in a comma-separated format that can be copied into a spreadsheet. The code used to generate total counts was written with the assistance of Omega, a Napari plugin that utilizes ChatGPT to process and analyze microscopy images [38]. We recommend Omega for assistance in writing code if users require a different output than the one our code generates.

**Figure 9:**
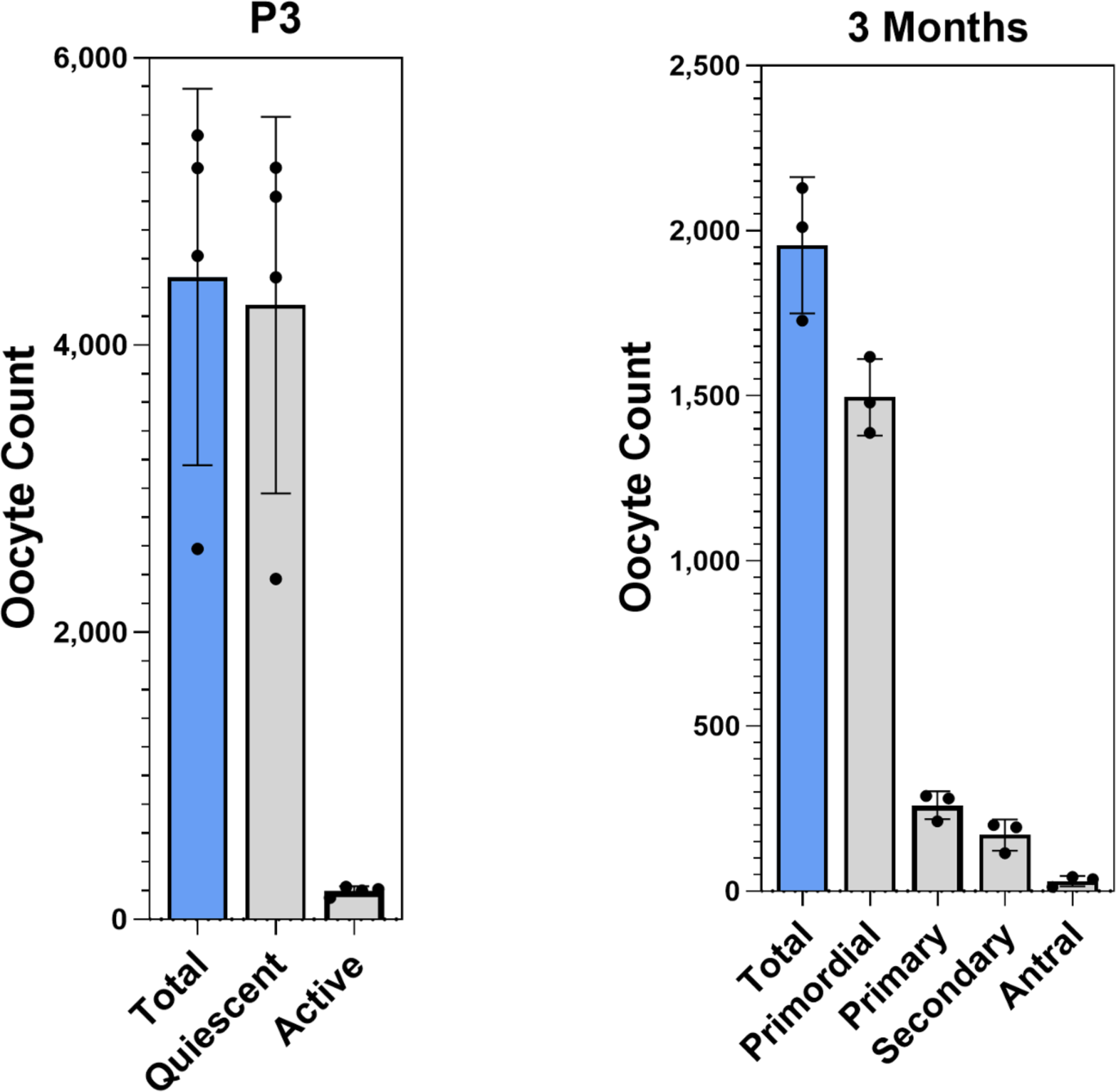
OoCount quantifies oocytes in 3D images of ovaries. A. Representative data from the OoCount workflow showing the total number of oocytes, the number of quiescent oocytes, and the number of active oocytes from four P3 ovaries. B. Representative data from the OoCount workflow showing the total number of oocytes and the number of oocytes at each growth stage in three 3-month-old ovaries.

### Creating a custom StarDist model

Because our model of StarDist was trained on images produced by our lab, we understand that our pipeline may not be versatile enough to segment oocytes from images with different features from our oocyte images. If the images generated in a different lab are incompatible, or do not yield satisfactory results with our versions of StarDist-OoCount, the following steps can be used to customize StarDist-OoCount. This process will require the user to create a new training dataset and retrain our StarDist model. This section uses tools explained in previous sections, so if needed, reference previous sections for additional information.

9.1.1 To train StarDist to segment oocytes from images unlike those produced in our lab, we recommend creating a new training dataset to retrain our StarDist-OoCount model. Create a folder to store the new training dataset. Within it, make two subfolders, one called “Images” and the other called “Masks.”

9.1.2 Begin by opening images as a .tif file in Napari. Click Plugins > Napari-StarDist. Run our model of StarDist following instructions from 7.2, remembering to use the model appropriate for the stage or image scale of ovaries being examined (adult or perinatal).

9.1.3 Once StarDist has run and segmented the oocytes, save the labels layer in “Masks’’ and the image layer in “Images.” Assuming that our model of StarDist is insufficiently or inaccurately segmenting oocytes in this image, the labels produced by StarDist will need to be corrected (see video 6, https://napari.org/stable/howtos/layers/labels.html). This can be done using any type of pen tablet, or a computer mouse. *To remove labels*, use the eyedropper tool to select a label that needs to be removed. Once selected, erase the label with the eraser tool, making sure it has been erased from all Z-stacks. *To edit pre-existing labels,* use the eyedropper tool to select a label that needs to be edited. Once selected, change the size/shape/depth of the label using the paintbrush tool. To add a new label, press “m” on the keyboard, and paint over the missing oocyte, making sure to paint in all Z slices where the oocyte is present. Repeat this for several images or regions of interest (ROI). **Note**: StarDist does not support sparse annotations, so it is essential to make sure that all oocytes in the image or ROI are labeled. Save the corrected labels layer in the “Masks” folder, making sure to name it exactly as its corresponding file in “Images”. The resulting image-mask pairs will constitute your training dataset. Make sure to create a copy of the directory for the StarDist-OoCount model (_Adult or _Perinatal) that you wish to retrain in the directory that contains “Images” and “Masks” folders. **Note**: for optimal training and accuracy, we recommend that your newly produced training data be scaled to match the training data we originally used to create the StarDist-oocount models. For this, we provide a python script (resample_tiffs.py) that automatically resamples x and y dimensions for all images and masks .tif files present within a specific directory. To run this script, copy the .py file into the directory where your images are located, and open a terminal from there. Run the script by typing “python resampleTiff.py’’ and press enter. The script will prompt you to set a rescaling factor. Our adult model expects an x and y value of 1.2μm and the perinatal model expects x and y dimensions of 0.603μm. Divide these values by your x and y values to determine the scaling factor to use. Resized images can now be used as StarDist training data.

9.2.1 To retrain our StarDist model, we recommend using DL4MicEverywhere [22]. This platform offers easy-to-use Jupyter notebooks that utilize CNNs to retrain StarDist. First, download Docker Desktop [39] (https://www.docker.com/products/docker-desktop/). Docker is a tool that allows the deployment of applications and all their dependencies as a single unit of software, referred to as a “container”. Open DockerDesktop and create an account. Now that Docker is installed, download the ZIP file of the DL4MicEverywhere Repository from Github [22] (https://github.com/HenriquesLab/DL4MicEverywhere/blob/main/docs/USER_GUIDE.md).

9.2.2 Open Docker Desktop. Within the downloaded DL4MicEverywhere file, double click the launcher with the appropriate operating system (Windows, MacOs, Linux). A window will open with options for notebooks to run (Figure 10A). Select DL4Mic notebooks and select the notebook for StarDist 3D. Enter the path for the folder containing the training dataset in “data”. Enter the path for the directory where you wish to store the newly trained model in “results”. Select Allow GPU and click Run. From here, the required packages will install and the StarDist 3D Jupyter notebook will open in a browser.

**Figure 10:**
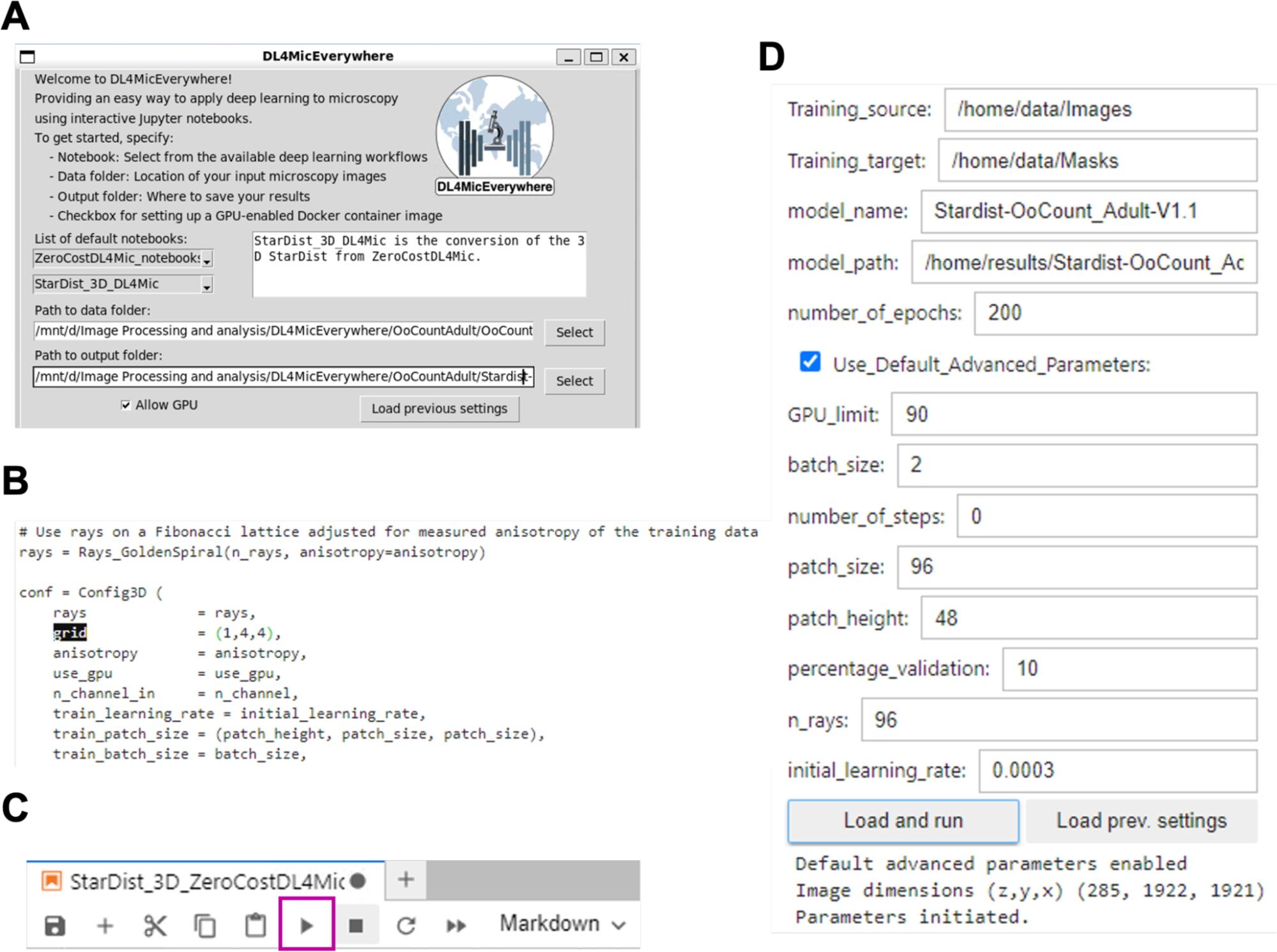
Retraining StarDist in DL4MicEverywhere. Screenshots from setting up the DL4MicEverywhere notebook to retrain StarDist. A. Once DL4MicEverywhere has been launched, a window will open with options for notebooks to run. Settings should look like those in this figure. B. Within the Jupyter notebook, the grid will need to be changed to “grid = (1,4,4).” C. The “Play” button can be found at the top of the notebook (highlighted here in a magenta box). D. When setting up the training, the settings of Section 3.1 of the Jupyter notebook should look similar to the settings above.

9.2.3 When prompted, select a Python kernel. While the code in the Jupyter notebook may be intimidating, there is only one line of code that must be modified. Use Ctrl/cmd+F to search for the word “grid.” Find the line of code where the grid is defined. Change “grid = grid” to “grid = (1,4,4)” (Figure 10B). This modification extends the field of view of the neural network, and allows segmentation of larger objects, such as large growing oocytes. Click play through the code chunks starting at 1.1 until section 3.1 of the StarDist 3D Jupyter notebook (Figure 10C, outlined in a magenta box).

9.2.4 At Section 3.1 of the StarDist 3D Jupyter notebook, in “training source”, enter the path for your training images; in “training target”, enter the path for your training masks (as seen in Figure 10D). The paths can be found by right-clicking the directories in the file browser in the left hand of the jupyter notebook window. Make sure to add the prefix /home/ to the path. These paths should typically look like: /home/data/Images; /home/data/Masks; /home/results/[name for your retrained model]. For “number of epochs”, enter 200 (Figure 10D). The rest of the settings should be left as default. Click load and run. This will display an example image-mask pair from the training dataset. At Section 3.2 of the notebook, don’t enable data augmentation. Click load and run. At Section 3.3 of the StarDist 3D Jupyter notebook, select “Use pretrained model” and use the dropdown bar to select Model from file. Copy the path to the OoCount-StarDist model you wish to retrain. Click “load and run”. Press play through both code chunks in Section 4 of the notebook. This will begin the training.

9.2.5 To evaluate the newly trained model, play the code chunks in Section 5. This can also be achieved by using the new model in StarDist-Napari on an image that was not a part of the training dataset.

### Troubleshooting

#### Choosing a classifier plugin

While we suggest using APOC to classify oocytes into growth stages, we understand that it may not be ideal because it has to be retrained and corrected for each batch of images. Training APOC is quick because it utilizes sparse annotation, which is a strength of this classifier. We also recommend using an activation marker, such as NR5A2, and to not rely only on the size of the oocyte to determine activation, as certain mutations or strain differences may affect the size of the oocyte independently from growth stage and yield suboptimal classification accuracy. If APOC does not yield satisfactory results or is becoming too cumbersome to train, we suggest trying another classifier Napari plugin called Svetlana [40] (https://pypi.org/project/napari-svetlana/). Svetlana leverages CNNs to create a reusable trained model for classification, meaning the user only has to train it once and the trained model can be used on new batches of images. Svetlana has a steeper learning curve, but some users may find it preferable for their workflow. For more information on Svetlana, we suggest reading the user guide (https://www.napari-hub.org/plugins/napari-svetlana). We also anticipate new classifiers will continue to be developed and made available as Napari plugins. The Napari-hub (https://www.Napari-hub.org/) catalogs all available Napari plugins, and is the ideal place to search for new plugin releases and plugin updates.

#### Optimizing DL4MicEverywhere

If retraining our StarDist model is necessary, it may require optimization beyond what is outlined in the section above. Some considerations when retraining StarDist include deciding how many epochs to run and the type of computing power available. If a training dataset is a set of flashcards that the neural network will learn from, the number of epochs reflects the number of times the neural network reviews the flashcards, or the training dataset. It may be intuitive to believe that more epochs would be better, however, overtraining the model can lead to overfitting and make it less effective. We suggest using somewhere between 100 and 200 epochs. **Note**: Increasing the number of epochs will also make the training longer and require more computational resources. If computational resources are a limitation, we recommend looking into computer clusters available through your institution or online computer clusters, such as google colab (https://colab.google/).

#### Resources for troubleshooting

Wherever possible, we have included the links to troubleshooting resources for using OoCount. All the programs included in OoCount have excellent documentation and tutorials, which have allowed us to learn these programs and incorporate them into OoCount. In addition to these program-specific resources, our lab will have additional resources on our website (www.mckeylab.com) for troubleshooting OoCount, with an option to contact us with questions.

## Conclusions

Ovarian follicle counts are a commonly used and insightful readout in ovarian biology research. With recent advances in 3D imaging, it is easier than ever to visualize the entire ovary and all the oocytes within it. However, analysis of these large images has been a major roadblock, as counting oocytes from these images is time consuming and prone to error. By leveraging machine learning tools, we have developed OoCount, an open-source, accessible workflow to quickly and efficiently segment oocytes and classify them based on growth stages in perinatal and adult mouse ovaries. Currently, OoCount successfully segments oocytes from 3D images of perinatal and adult mouse ovaries and yields results similar to previously published studies [4,41–44]. However, it is important to note that OoCount was developed and run on CD-1 mice and it has been documented that there are differences in oocyte quantity based on mouse strain [5]. While OoCount is not the first computational workflow for oocyte segmentation and classification [11,16,45], OoCount is advantageous because it utilizes open-source resources, and relies on an oocyte activation marker for classification, allowing for efficient, accessible oocyte segmentation and classification. While OoCount was optimized for mouse tissue and we did not test OoCount on ovaries from other animals, we expect it to be easily transferable to other species with some retraining.

Here, we detailed an end-to-end pipeline consisting of protocols optimized in our lab and workflows to segment and classify oocytes from 3D images of perinatal and adult mouse ovaries. While we hope that OoCount will become a widely adopted tool, we understand there are limitations. Follicular atresia is not accounted for in this pipeline. Any atretic follicles are classified along with other follicles or not segmented due to low DDX4 signal. Atretic follicles that are segmented by StarDist could be classified in APOC if the user has used an antibody to act as a marker for atresia. Additionally, APOC is limited in its ability to automatically distinguish between secondary and early antral follicles in the adult ovary based only on the expression of DDX4 and NR5A2. We highly recommend careful inspection and manual correction of the APOC predictions for optimal count accuracy. For users particularly interested in distinguishing between these stages, we recommend using a stage-specific marker for better classification in APOC. The entire workflow was generated using a PC with a Windows operating system and may require additional troubleshooting if using a different operating system. We have included links to resources for troubleshooting the different software packages within this workflow (Table S3). As of now, our classification uses APOC, which is the most user-friendly classifier we encountered, but APOC may require the user to retrain it for each set of images. In the future, we hope to develop our own classifier, specific to classifying oocytes in 3D images of mouse ovaries. In addition, we plan to create a plugin or software containing the entire pipeline. We hope this will increase the ease of use of OoCount and provide the field with a much needed standardized tool for counting oocytes in 3D images of mouse ovaries.

## Acknowledgments

The authors would like to thank all members of the Capel Lab, McKey Lab, and Roberson Lab for helpful discussions and feedback during the development of this project. The authors would also like to thank Victor A. Ruthig and Elle C. Roberson for feedback on the initial drafts of the manuscript. Special thanks to Leandro Lovisolo for essential training and guidance on computational aspects of this study, and to Wendy Zhang for testing the protocol. We would also like to acknowledge Sofia Batchvarova, who patiently annotated an entire 3D mouse ovary in the first iteration of this project.

## Author contributions

JM and BC conceptualized the project. LF, CB and JM designed and optimized the computational pipeline. LF, ASM and JM performed optimization of the bench protocols and data collection. LF produced the initial draft of the manuscript and figures. BC, CB, ASM and JM edited the manuscript. JM provided funding for the project.

## Video captions

**Video 1:** Fly-through of Z-stacks of a perinatal (P3) mouse ovary immunostained with DDX4 (oocytes) and NR5A2 (activated oocytes and granulosa cells).

**Video 2:** Fly-through of Z-stacks of an adult (3 month) mouse ovary immunostained with DDX4 (oocytes) and NR5A2 (activated granulosa cells).

**Video 3:** Screen-captured video of steps for opening Napari from Anaconda Navigator.

**Video 4:** Screen-captured video showing basics for navigating the Napari interface.

**Video 5:** Screen-captured video showing how to run StarDist-OoCount on a P3 mouse ovary in Napari.

**Video 6:** Screen-captured video showing how to correct mis-segmented oocytes from StarDist-OoCount in Napari.

**Video 7:** Screen-captured video showing how to annotate a P3 mouse ovary for classification using APOC in Napari.

**Video 8:** Screen-captured video showing how to train APOC to automatically classify oocytes from annotations.

**Video 9:** Screen-captured video showing how to correct APOC results.

**Video 10**: Screen-captured video showing how to use the provided code to generate final oocyte counts.

## Supplemental Figures

**Supplemental Figure 1:**
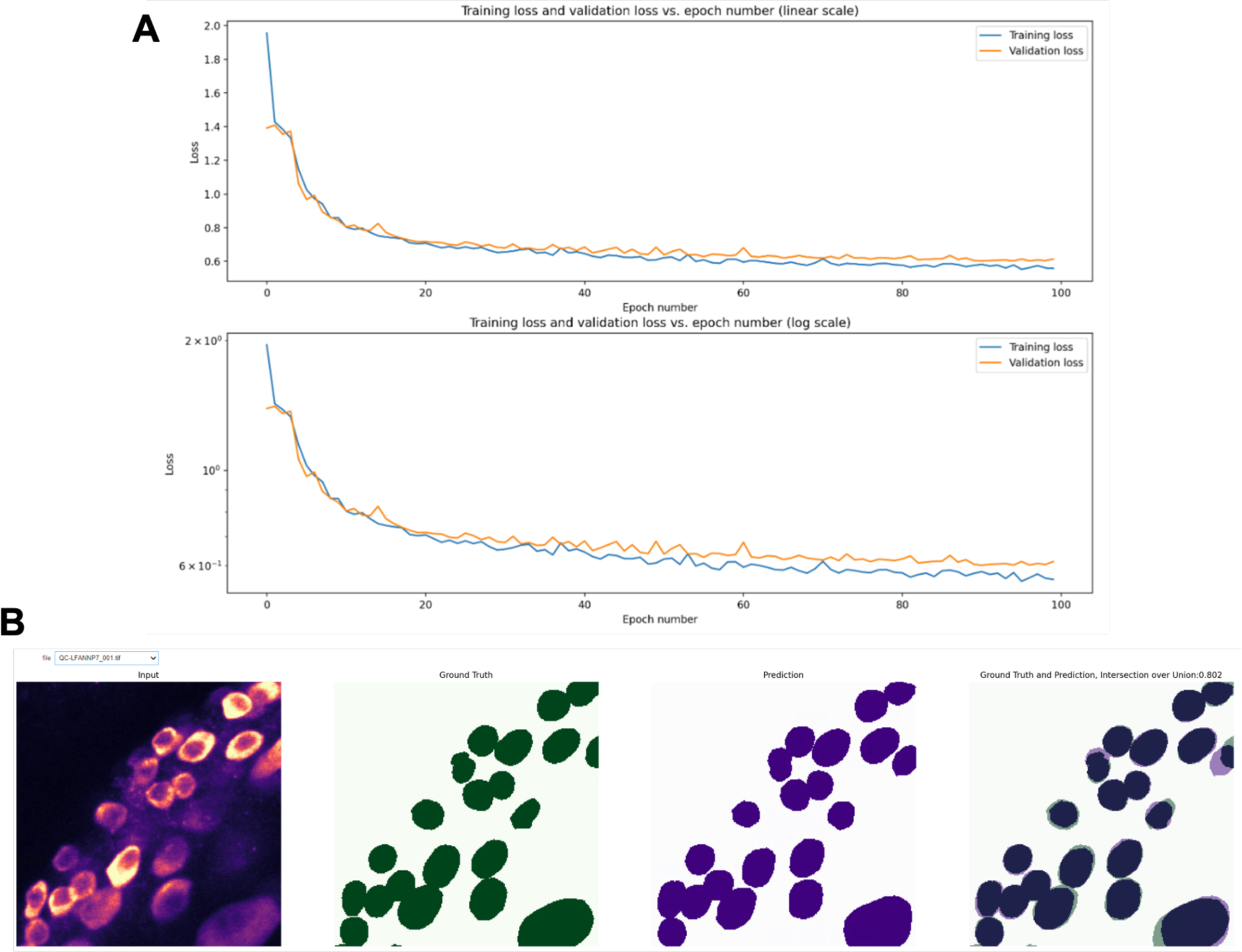
Validating training efficacy. A. Training loss and validation loss plot generated by the DL4MicEverywhere StarDist 3D training notebook. B. Intersection over Union prediction generated by DL4MicEverywhere StarDist 3D training notebook.

**Supplemental Figure 2:**
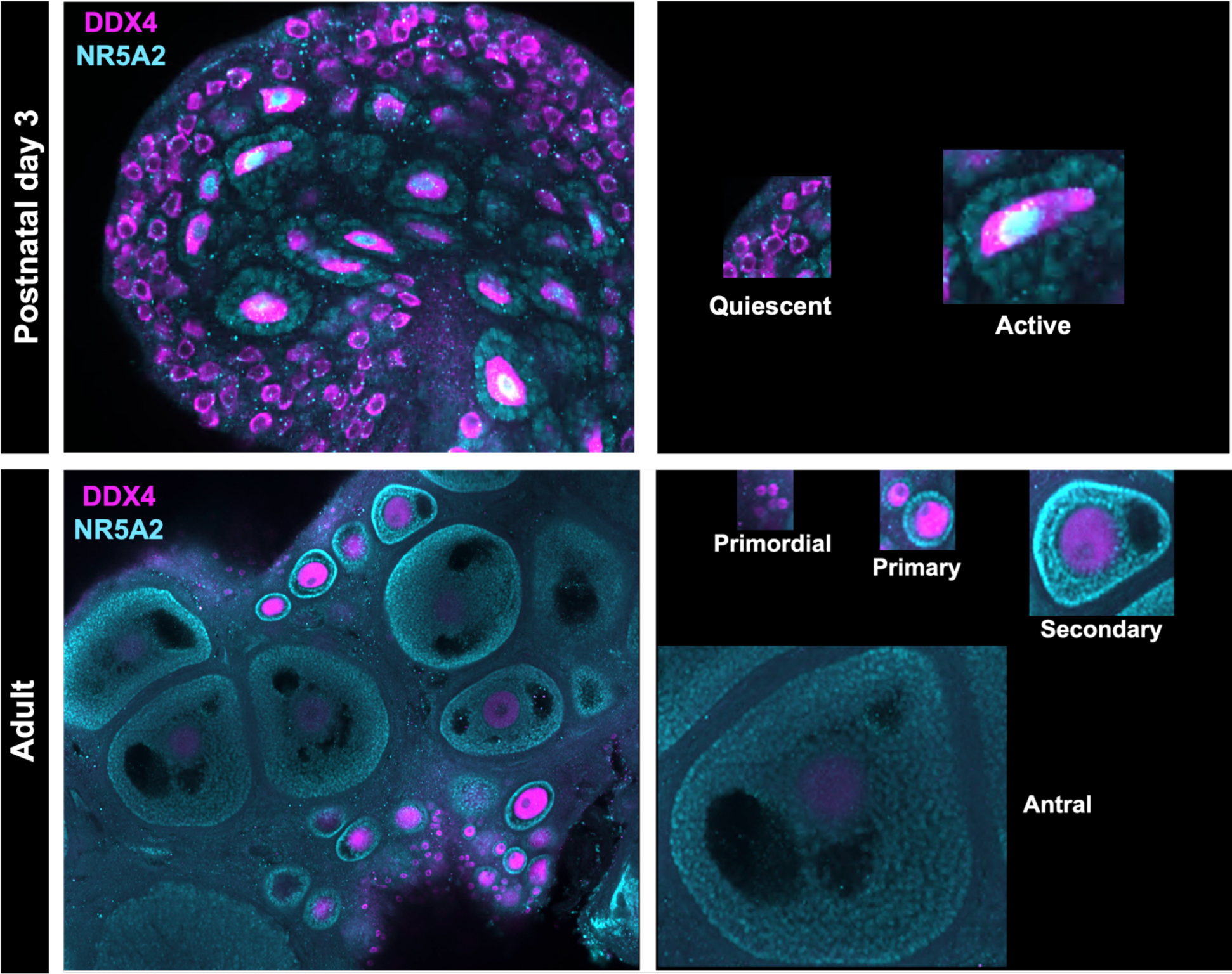
Determining growth stage of oocytes in perinatal and adult ovaries. In perinatal ovaries, DDX4 is expressed in all oocytes and NR5A2 is expressed in the nuclei and granulosa cells of active oocytes. Oocytes are classified as quiescent if NR5A2 is not expressed and active if NR5A2 is expressed. In adult ovaries, DDX4 is expressed in all oocytes and NR5A2 is expressed in the granulosa cells of active oocytes. Oocytes are classified as: primordial if there are no NR5A2+ granulosa cells surrounding the oocyte, primary if there is one layer of NR5A2+ granulosa cells surrounding the oocytes, secondary if there are multiple layers of NR5A2+ granulosa cells surrounding the oocytes, and antral if there are multiple layers of NR5A2+ granulosa cells surrounding the oocytes and a distinguishable antral cavity.

## Supplemental Tables

**Supplemental Table 1.** List of antibodies used in this study.

| Primary Antibody | Host Species | Dilution | Supplier | Catalog Number | RRID | Secondary Antibody | Dilution | Supplier | Catalog Number | RRID |
| --- | --- | --- | --- | --- | --- | --- | --- | --- | --- | --- |
| DDX4/MVH | Rabbit | 1:250 | abcam | ab13840 | AB_443012 | Alexa Fluor 647 Donkey Anti-Rabbit | 1:500 | Jackson ImmunoResearch | 711605152 | AB_2492288 |
| GDF9 | Rabbit | 1:250 | Proteintech | 29309-1-AP | AB_3086112 |  |  |  |  |  |
| Human VASA (DDX4) | Goat | 1:250 | R&D Systems | AF2030 | AB_2277369 | Cy3 anti-goat | 1:500 | Jackson ImmunoResearch | 705165147 | AB_2307351 |
| NR5A2 | Goat | 1:500 | abcam | ab223211 | AB_3096000 |  |  |  |  |  |

**Supplemental Table 2.** Materials recommended for mounting perinatal and adult ovaries for in toto imaging.

| **Fetal and Perinatal Whole Mount Immunofluorescence** | **Source** | **Catalog Number** |
| --- | --- | --- |
| 40mm Cover Slips | Fisher Scientific | 12-541-021 |
| Sally Hansen Hard as Nails | Drugstore | N/A |
| 3D Printed Slide | NIH 3D Exchange | https://3d.nih.gov/builds/550 |
| 60mL Petri Dish | Fisher Scientific | 08-747A |
| Dumont #5 Forceps | Fine Science Tools | 11252-20 |
| Argos Technologies Syringe Filter, .22um, PP/PVDF | Cole Parmer | 04396-32 |
| Fisherbrand Sterile Syringes for Single Use | Fisher Scientific | 14-955-464 |

| **Adult Whole Mount Immunofluorescence** | **Source** | **Catalog Number** |
| --- | --- | --- |
| 40mm Cover Slips | Fisher Scientific | 12-541-021 |
| 3D printed slide | NIH 3D Exchange | https://3d.nih.gov/builds/550 |
| Silicon sealant or superglue | Drugstore | N/A |
| Dumont #5 Forceps | Fine Science Tools | 11252-20 |
| UV Flashlight LED | VWR | 470311-976 |
| Argos Technologies Syringe Filter, .22um, PP/PVDF | Cole Parmer | 04396-32 |
| Fisherbrand single-use sterile Syringes | Fisher Scientific | 14-955-464 |

**Supplemental Table 3.**
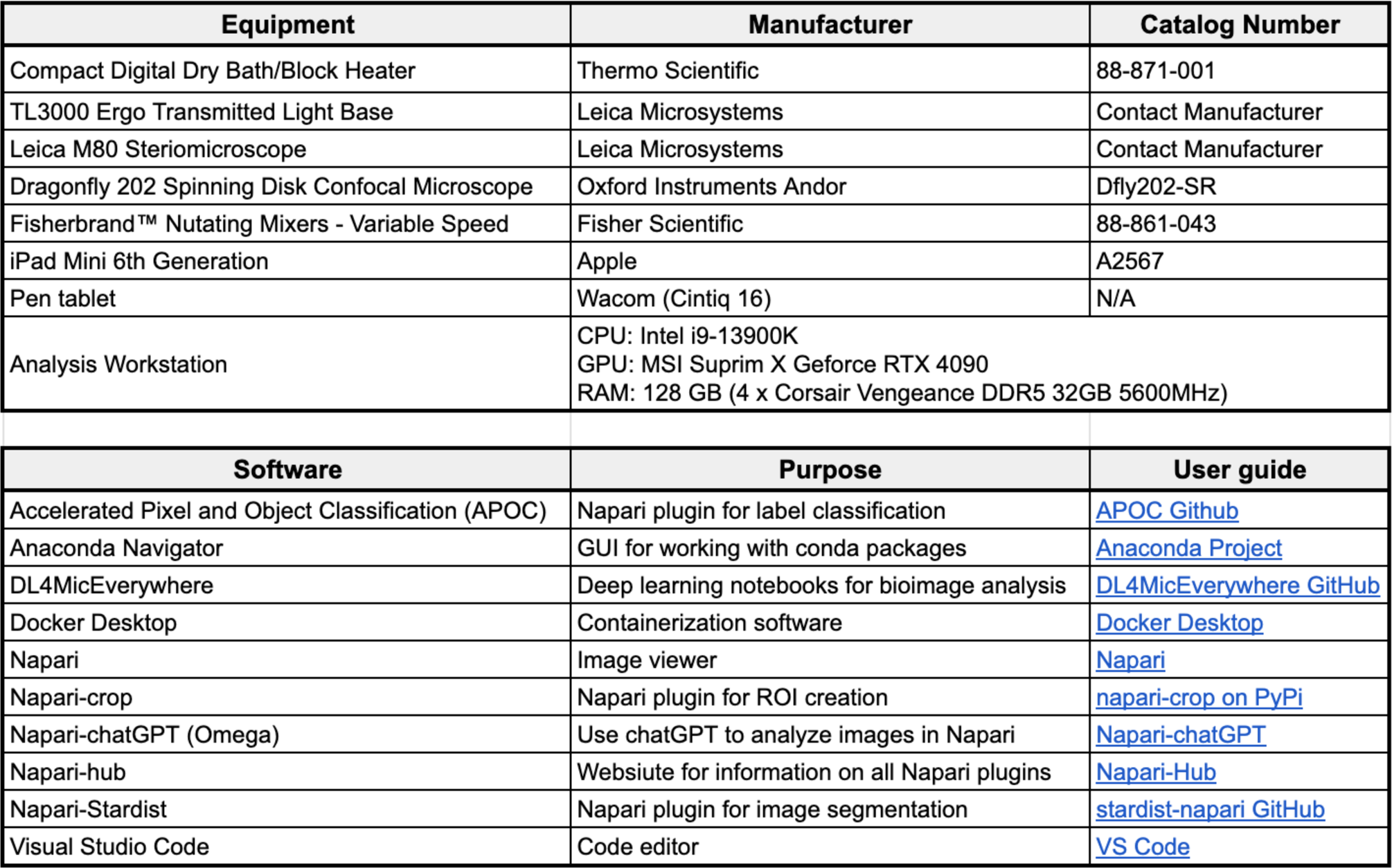
Equipment and software used to develop and apply the OoCount pipeline.

